# Functional characterisation and identification of a core interactome of *Plasmodium* regulatory protein Alba4

**DOI:** 10.64898/2026.11.10.765626

**Authors:** Meghan E. Zadow, Ghizal Siddiqui, Balu Balan, Joshua P. Morrow, Aaron R. Jex, Christopher A. MacRaild, Darren J. Creek, Danny W. Wilson

**Affiliations:** Reseach Centre for Infectious Diseases, School of Biological Sciences, Adelaide University, Adelaide 5005, Australia; Institute for Photonics and Advanced Sensing (IPAS), Adelaide University, Adelaide 5005, Australia; Drug Delivery, Disposition and Dynamics, Monash Institute of Pharmaceutical Sciences, Monash University, Parkville, Victoria 3052, Australia; The Walter and Eliza Hall Institute of Medical Research, Parkville, Australia; University of Melbourne, Melbourne, Australia; Burnet Institute, Melbourne 3004, Victoria, Australia

## Abstract

*Plasmodium* spp. parasites have complex lifecycles that rely on strict regulation of gene and protein expression. Current evidence suggests that *P. falciparum* has a relatively limited repertoire of canonical transcriptional regulators and RNA interference (RNAi)-associated small RNAs compared to model eukaryotes, potentially reflecting a greater reliance on RNA binding protein (RBP)-mediated post-transcriptional regulation (PTR). Among these RBPs, the acetylation lowers binding affinity (Alba) family of DNA/RNA-binding proteins has been implicated in RNA metabolism and regulation of mRNA stability and translation. In *Plasmodium* spp. parasites, the Alba family has expanded to include up to six members, with sequence divergence and structural modifications beyond the conserved Alba domain leading to diversification of family members, some of which appear restricted to *Plasmodium* and related parasites. In this study, we were unable to truncate PfAlba4, but successfully generated an endogenously tagged PfAlba4 parasite line, representing the first successful instance of endogenous tagging of a PfAlba family protein. Utilising this transgenic line, we confirm ubiquitous expression throughout blood stage development and its predominantly cytosolic localisation. Conditional loss of PfAlba4 through the knock-sideways approach demonstrated an important role for PfAlba4 in blood stage development and proteomics analysis defined a protein interaction network of 67 significantly enriched proteins, with notable enrichment of proteins potentially involved in RNA processing and regulation. Further characterisation of one of these proteins, the uncharacterised protein PF3D7_0706500, revealed a partially overlapping network of proximal proteins enriched for RNA-regulatory factors, suggesting potentially shared roles in PTR. RNA Immunoprecipitation-Sequencing (RIP-seq) identified 692 PfAlba4 associated transcripts, with an enrichment of a pyrimidine-rich motif in their 3’ untranslated regions, suggesting a potential sequence-dependent mechanism of RNA recognition. Integration of proteomic and RIP-Seq datasets in late blood stage parasites revealed a consistent association with RNA processing and regulatory pathways, implicating PfAlba4 in post-transcriptional regulation. Together these findings establish PfAlba4 as an important component of the RNA-regulatory landscape in *P. falciparum* and highlight the importance of PfAlba proteins for malaria parasite blood stage development.

## Introduction

Malaria is a persistent mosquito-borne disease caused by *Plasmodium* spp. parasites. In 2024, there were ∼282 million cases of malaria resulting in ∼610,000 deaths (WHO, 2025). While there are multiple *Plasmodium* species known to infect and cause disease in humans, the majority of cases and severe disease are attributable to the species *Plasmodium falciparum*. Despite continued efforts towards eradication, malaria remains endemic in 80 countries with the WHO sub-Saharan African region shouldering the largest burden of cases (WHO, 2025). Effective control and treatment of *P. falciparum* malaria parasites is hampered by their remarkable ability to adapt and develop resistance to available therapeutics, as well as their evolution of genetic mutations that allow them to evade detection by the most commonly used rapid diagnostic tests (Kong *et al*, 2021).

Human *P. falciparum* infection begins in the liver before progressing to the blood stage of infection, which is the stage at which symptomatic disease occurs (Figure 1a). Immediately after invading a red blood cell (RBC), the parasite modifies the host cell and undergoes a period of growth and nutrient acquisition termed the ring stage. This is followed by the trophozoite stage, during which the single-nucleated parasite begins replicating its DNA before transitioning to the multinucleated schizont stage. The parasite continues to asexually replicate its DNA until ∼48-hours post-invasion (h.p.i.) when it then ruptures the host cell to release ∼16-32 progeny into the bloodstream to continue the infection cycle. Invasion of a new RBC is rapid, thus limiting the parasite’s exposure to host immune defences. During the blood stage a small proportion of parasites undergo sexual commitment to form male and female gametocytes. Female and male gametocytes are necessary for transmission from the human host to the mosquito vector and enable sexual reproduction to take place in the mosquito.

**Figure 1.**
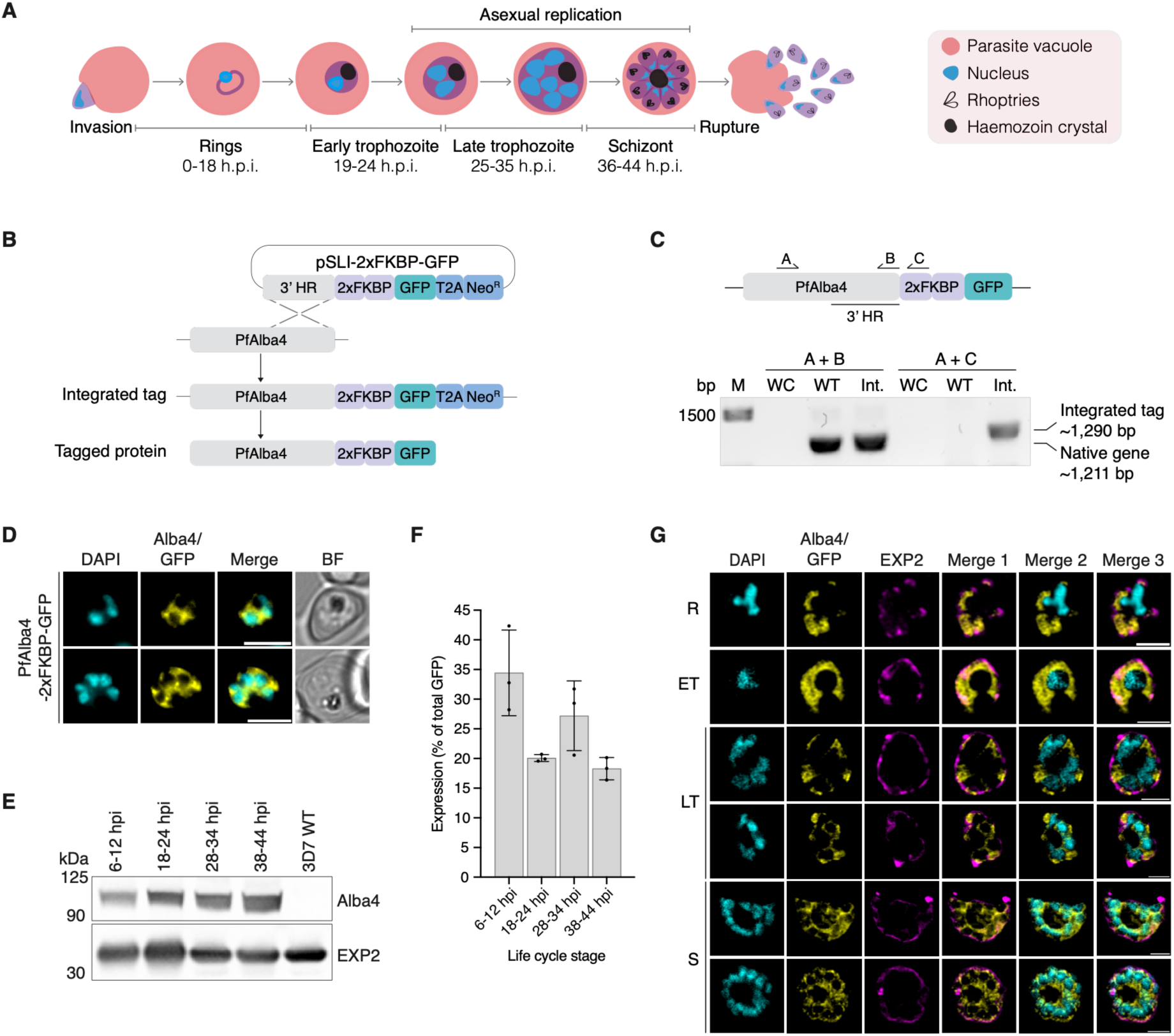
Endogenous tagging and characterisation of PfAlba4. **A)** *Plasmodium falciparum* blood stage development is considered to adhere to a ∼48-hour rupture/invasion cycle. Following parasite invasion of a red blood cell, the parasite progresses through a period of growth and nutrient acquisition (ring stage) before commencing replication (trophozoite stage) and undergoing asynchronous, asexual replication in a process termed ‘schizogony’ (schizont stage). The lab-adapted 3D7 wildtype strain used in this study follows a ∼44-hour cycle. **B)** C-terminal endogenous tagging of *Pfalba4* was achieved by selection linked integration (SLI) (Birnbaum *et al*, 2017). A vector plasmid containing a 3’ homology region (HR) for *Pfalba4* followed by a 2xFKBP-GFP tag, skip peptide (T2A) and a neomycin resistance cassette was transfected into 3D7 wildtype parasites, facilitating single crossover recombination and tag integration (PfAlba4^2xFKBP-GFP^). **C)** Successful tag integration was confirmed by PCR. Primers A and B correspond to a 1,211 bp region of the native gene. Bands of expected size were observed for both the 3D7 wildtype (WT) and PfAlba4^2xFKBP-GFP^ (Int.), no band was observed in the water control (WC). Primer A targets a region of *Pfalba4* upstream from the 3’ HR, and C corresponds to a region in the FKBP tag. Successful integration is indicated by the presence of a ∼1,290 bp band present in the PfAlba4^2xFKBP-GFP^ (Int.) lane only. **D)** Live cell microscopy confirms expression of endogenous tag in the PfAlba4^2xFKBP-GFP^ line. The GFP signal in yellow indicates a cytoplasmic localisation for PfAlba4. Nuclei were stained with DAPI (cyan). Scale bars = 5 μm. **E)** Western blot analysis shows ubiquitous expression of PfAlba4 across the blood stage and, **F)** quantification of protein expression across n=3 biological replicates for the PfAlba4^2xFKBP-GFP^ line. Tagged PfAlba4 (∼100 kDa) was detected by anti-GFP and anti-EXP2 (∼33 kDa) was used as a loading control. Individual data points are represented as well as the mean (columns) and standard deviation (error bars). **G)** Immunofluorescence microscopy showing the cytoplasmic expression pattern of PfAlba4 in rings (R), early trophozoites (ET), late trophozoites (LT) and schizonts (S) in the PfAlba4^2xFKBP-GFP^ line. Nuclei are represented in cyan, PfAlba4 in yellow, and the parasite vacuole protein PfEXP2 in magenta. Merge 1 – PfAlba4 and PfEXP2, Merge 2 – DAPI and PfAlba4, Merge 3 – DAPI, PfAlba4, and PfEXP2. Scale bars = 2 μm.

The complexity of this life cycle, spanning the mosquito vector and human host, infection of multiple tissue types, and utilisation of both sexual and asexual methods of DNA reproduction, necessitates constant adaptation by the parasite to harsh and changing environments, varying metabolic demands, nutrient availability, and host immune responses. Such adaptations would require a high degree of gene and protein regulation. However, the mechanisms by which the parasite achieves this regulation remain poorly understood.

Many model organisms rely heavily on transcriptional regulation in order to coordinate gene and protein expression. For example, in humans, gene expression is heavily governed by a suite of proteins termed ‘transcription factors’ (Lambert *et al*, 2018). In *P. falciparum*, a relatively limited repertoire of canonical transcription factors has been identified compared to model organisms to date (Balaji *et al*, 2005; Gardner *et al*, 2002; Painter *et al*, 2011; Templeton *et al*, 2004). It has also been observed that transcript abundance poorly correlates with protein abundance in these parasites, with ∼30% of transcripts reported to accumulate hours before their corresponding proteins were detected, highlighting the likely importance of post-transcriptional regulation in these parasites (Foth *et al*, 2011; Foth *et al*, 2008; Le Roch *et al*, 2004).

RNA-binding proteins (RBPs) are a very large and abundant functional group of proteins with known involvement in post-transcriptional regulation. Notably, *Plasmodium* spp. parasites encode a larger repertoire of RBPs in their genomes than transcription factors, suggesting an important contribution of RBP-mediated post-transcriptional regulation (Reddy *et al*, 2015). One family of RBPs that appear to have expanded in *P. falciparum* are the Acetylation lowers binding affinity (ALBA) domain containing protein family, of which six members have so far been identified *(Chêne et al, 2012; Reddy et al., 2015)*. Alba proteins are conserved across all three domains of life, although the number of family members varies considerably among organisms. For example, humans encode just 2 of these proteins (Rpp20 and Rpp25) while the allotetraploid cotton species *G. hirsutum* encode 33 Alba domain-containing proteins (Guerrier-Takada *et al*, 2002; Li & Altman, 2001; Magwanga *et al*, 2019). Alba proteins are reported to exhibit considerable functional diversity, including roles in critical cellular processes such as stress responses, virulence gene regulation, and translational regulation in protozoan parasites (Zadow *et al*, 2025).

Our understanding of the functional roles of *Plasmodium* Alba proteins is limited, particularly in the major human malaria parasite *P. falciparum*. Here, we focus on PfAlba4, previously identified as essential to *P. falciparum* blood stage growth in a genome-wide saturation mutagenesis study (Zhang *et al*, 2018). This indispensability is supported by studies reporting that PfAlba4 is refractory to endogenous tagging, leading to a reliance on episomal overexpression systems and in-house antibodies for functional characterisation to date (Acharya *et al*, 2025; Chêne *et al*., 2012; Vembar *et al*, 2015). It is notable however that the orthologous PyAlba4 from the rodent-infecting *P. yoelii* has been reported to be dispensable for asexual blood stage development, evidenced by the successful generation of PyAlba4^null^ parasites (Muñoz *et al*, 2017; Walker & Lindner, 2019). These contrasting findings highlight the need for further functional characterisation of PfAlba4 to clarify its contributions to *P. falciparum* blood-stage development.

In this study, we successfully generated an endogenously tagged PfAlba4 transgenic line which, to our knowledge, is the first successful instance of an endogenously tagged Alba protein in *P. falciparum*. Using this line, we investigated the role of PfAlba4 during *P. falciparum* asexual blood stage growth and characterised its associated protein and RNA interactions through complementary proteomic and transcriptomic approaches, providing new insights into its potential role in malaria parasite post-transcriptional regulation.

## Results

### PfAlba4 is ubiquitously expressed and localised to the cytoplasm during *P. falciparum*blood stage development

PfAlba4 is the largest of the PfAlba proteins at 42 kDa and consists of an N-terminal Alba domain and a C-terminal mitochondrial association factor 1 (Maf1)-like domain. PfAlba4 was previously assigned to the Rpp25-like group of Alba domain proteins, however, current evidence supports grouping of this protein lineage into an apicomplexan-specific PfAlba4-like cluster (Aravind *et al*, 2003; Jagadeesh & Vembar, 2024; Zadow *et al*., 2025). Previous studies of PfAlba4 in *P. falciparum* and its orthologues in the rodent-malaria parasites, *P. berghei* and *P. yoelii*, have implicated Alba4 proteins in several cellular processes including virulence gene regulation, stage differentiation, and stress responses (Acharya *et al*., 2025; Chêne *et al*., 2012; Mair *et al*, 2010; Muñoz *et al*., 2017).

To characterise the function of PfAlba4 during *P. falciparum* blood stage development, we generated a parasite line expressing PfAlba4 with an endogenous 2xFKBP-GFP tag to the C-terminus of 3D7 PfAlba4 (PfAlba4^2xFKBP-GFP^) using selection-linked integration (Figure 1b) (Birnbaum *et al*, 2017). Following generation of viable parasites, integration of the tag was confirmed by PCR and GFP expression of the tagged protein by live cell microscopy (Figures 1c, d; Supplementary figure 1a). To observe PfAlba4 expression across blood stage development, we analysed protein lysates harvested from four distinct time points, with western blot analysis indicating that this protein is ubiquitously expressed across blood stage development (Figure 1e, f; Supplementary data 1). Live-cell microscopy and immunofluorescence assays revealed predominantly cytoplasmic localisation of PfAlba4 across blood stage development, with apparent accumulation near the nuclear envelope and parasite plasma membrane during the schizont stage. These observations are consistent with previous reports of Alba4 localisation in *P. falciparum, P. berghei,* and *P. yoelii* (Figure 1g; Supplementary figure 1b) (Acharya *et al*., 2025; Chêne *et al*., 2012; Mair *et al*., 2010; Muñoz *et al*., 2017).

### PfAlba4 and its Maf1-like domain are important for blood stage development

Previous genome-wide saturation mutagenesis studies and unsuccessful attempts to tag PfAlba4, have suggested that PfAlba4 is essential for *P. falciparum* blood stage growth. In contrast, successful deletion of orthologous PyAlba4 in *P. yoelii,* suggest that Alba4 is dispensable for parasite growth in this rodent malaria (Muñoz *et al*., 2017; Walker & Lindner, 2019). Given these conflicting data, we next utilised two alternative approaches to investigate the functional importance of PfAlba4 during parasite blood stage development.

To determine whether the C-terminal Maf1-like domain of PfAlba4 is dispensable during the blood stage, we utilised a targeted gene deletion method (pSLI-TGD) and attempted to generate a parasite line where the Maf1-like domain is replaced by a GFP tag (PfAlba4^ΔMaf1-GFP^, Figure 2a; Supplementary figure 2a) (Birnbaum *et al*., 2017). Despite four independent attempts to replace the Maf1-like domain with a GFP tag, no viable parasites were observed after selection for integration (Figure 2a). This indicates that PfAlba4’s c-terminus and Maf1-like domain is essential for *P. falciparum* blood stage growth, supporting the importance of this protein for *P. falciparum* blood stage development.

**Figure 2.**
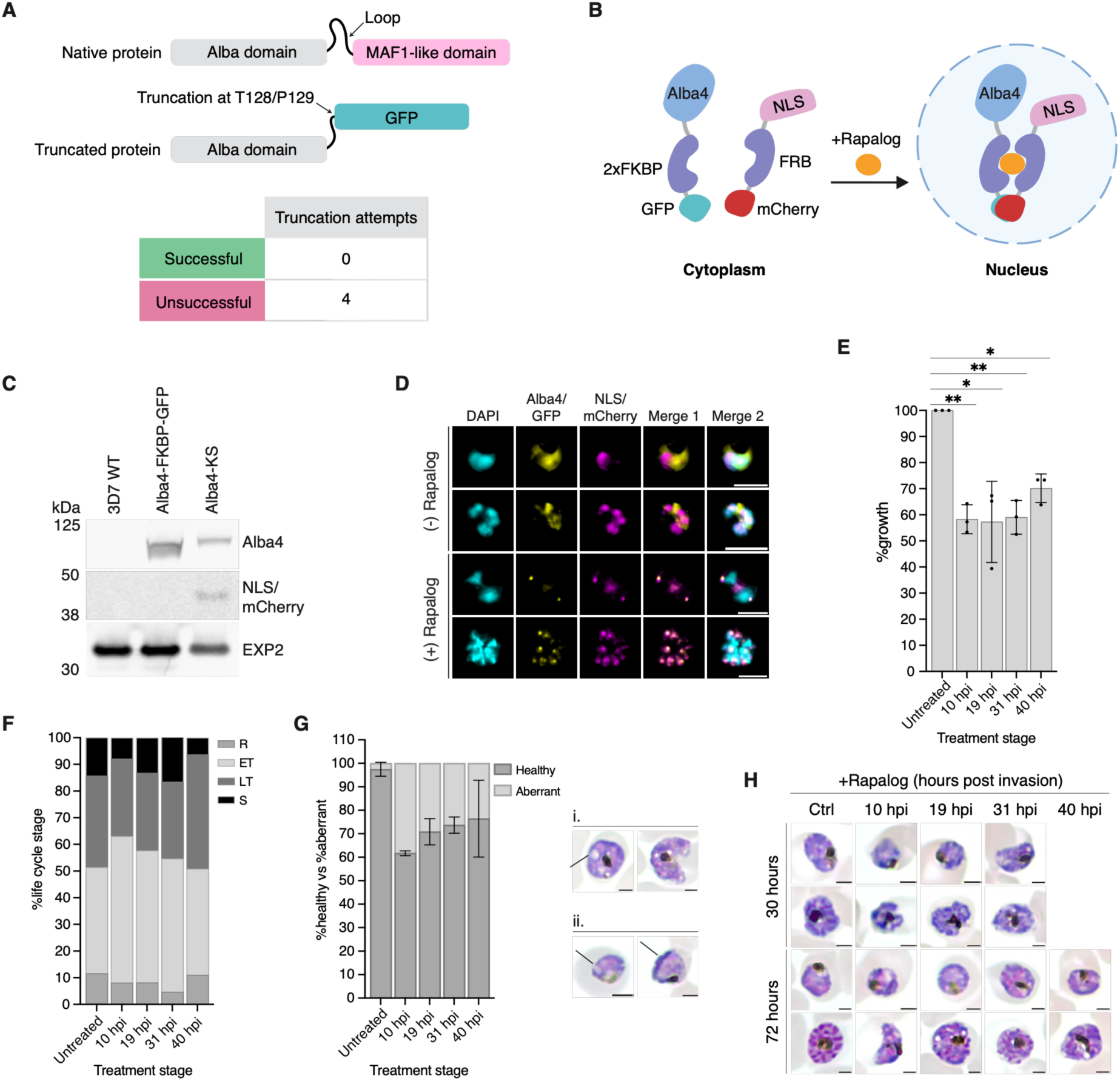
Phenotypic analysis of PfAlba4 during blood stage development. **A)** Removal of the PfAlba4 Maf1-like domain was attempted using targeted gene deletion (TGD)-SLI. A TGD transfection plasmid was generated with a 3’ HR for the Alba domain (52-384 nt) which would truncate PfAlba4 at T128 and insert a GFP tag in place of the Maf1-like domain. Four independent transfections failed to generate a viable PfAlba4^ΔMaf1-GFP^ transgenic line. **B)** Schematic representation of targeted and inducible protein mislocalisation. A nuclear mislocalising plasmid is co-transfected into PfAlba4^2xFKBP-GFP^ parasites (PfAlba4^2xFKBP-GFP/NLS-FRB-mCh^). Addition of the drug rapalog facilitates conditional dimerisation of the episomally expressed FRB subunit with the FKBP subunits appended to PfAlba4. The nuclear localisation signal (NLS) facilitates removal of PfAlba4 from the cytoplasm and its site of likely function to the nucleus. **C)** Western blots show retention of the nuclear mislocalising episome in the PfAlba4 transgenic line. Membranes were probed with anti-GFP to detect PfAlba4 (∼100 kDa), anti-mCherry to detect the mislocalisation episome (∼44 kDa), and anti-EXP2 (∼33 kDa) was used as a loading control. **D)** Live cell microscopy demonstrates differential mislocalisation of PfAlba4 in the presence or absence of rapalog for the PfAlba4^2xFKBP-GFP/NLS-FRB-mCh^ line. PfAlba4 (GFP in yellow) localises to the cytoplasm in the absence of rapalog and the mislocalising episome (mCherry in magenta) co-localises with the nuclei (DAPI in cyan). When rapalog is added the GFP and mCherry signals overlap and co-locate at the nuclear periphery. Merge 1 – GFP and mCherry, merge 2 – DAPI, GFP, mCherry. Scale bar = 5 µm, n=2 biological replicates**. E)** Nuclear mislocalisation of PfAlba4 negatively impacts parasite growth across the blood stage. Highly synchronised PfAlba4^2xFKBP-GFP/NLS-FRB-mCh^ parasites were treated with rapalog at pre-defined time points across the blood stage (10, 19, 31, and 40 h.p.i.) to induce mislocalisation of PfAlba4. Drug was removed at 48-hours post assay set-up and growth assessed at 72-hours. Growth is expressed as a percentage compared to the untreated control and is an average of n=3 biological replicates. Significance determined by paired two-tailed t-tests (\**p* <0.05, \*\**p* <0.01). Error bars = standard deviation. **F)** PfAlba4 mislocalisation in PfAlba4^2xFKBP-GFP/NLS-FRB-mCh^ parasites did not significantly impact stage transition however, **G)** mislocalisation resulted in a significant increase in aberrant phenotypes largely characterised by the presence of stress vacuoles and cellular disorganisation (i) and membrane abnormalities (ii). Error bars show standard deviation. **H)** Representative Giemsa images from each treatment group at 30- and 72-hours post assay set-up. Scale bars = 2 µm.

We next investigated the impact of mislocalisation of PfAlba4 away from its typical cytoplasmic localisation and to the nucleus using the knock-sideways approach (KS, (Birnbaum *et al*., 2017)). To achieve PfAlba4 mislocalisation, the endogenously tagged PfAlba4^2xFKBP-GFP^ parasite line was co-transfected with the plasmid encoding an FRB domain fused to a nuclear localisation signal (NLS-FRB-mCherry), enabling rapalog-dependent recruitment of PfAlba4 to the nucleus (Figure 2b). Episome retention for the resulting PfAlba4^2xFKBP-GFP/NLS-FRB-mCh^ line was confirmed by western blot (Figure 2c) and rapalog-induced mislocalisation of PfAlba4 (PfAlba4-KS) was verified by live-cell microscopy (Figure 2d; Supplementary figure 2b, c). Rapalog-induced PfAlba4 mislocalisation was commenced at the following developmental stages – 10 h.p.i. (rings), 19 h.p.i. (early trophozoites), 31 h.p.i. (late trophozoites), and 40 h.p.i. (schizonts). Rapalog was removed 48-hours post-assay set-up and parasite growth assessed at 72-hours alongside untreated PfAlba4-KS control parasites. Utilising this approach, we observed up to a 40% reduction in parasite growth across the blood stage for PfAlba4-KS compared to the untreated PfAlba4-KS control (Figure 2e; Supplementary data 1). To determine whether this growth impact was associated with delayed or stalled progression through blood stage development, we analysed Giemsa-stained blood smears, which revealed no significant differences in the distribution of life stages between the untreated and rapalog-treated PfAlba4-KS parasites (Figure 2f; Supplementary figure 2d). Given that the growth defect did not appear to be a result of delayed development, we next assessed any morphological differences between PfAlba4-KS untreated and rapalog-treated parasites. It was observed that all PfAlba4-KS treatment groups had an increased proportion of aberrant cell types compared to the untreated control parasites (Figure 2g, h; Supplementary figure 2e). This was largely characterised by the presence of stress vacuoles and membrane irregularities such as invagination and was most evident in parasites treated at ring stages where ∼40% of PfAlba4-KS cells displayed an aberrant morphology compared to ∼3% in the PfAlba4-KS untreated control parasites. Importantly, rapalog treatment of 3D7 WT parasites did not significantly affect parasite growth or morphology, indicating that the observed phenotypes were not attributable to rapalog treatment alone (Supplementary figure 3a-d).

Together, the unsuccessful attempts to truncate PfAlba4 and the growth and morphological defects observed following nuclear mislocalisation, indicate that PfAlba4 has an important role during parasite blood stage development.

### miniTurboID mediated proximity labelling identifies a PfAlba4-associated protein network enriched in RNA regulatory factors in late blood stage development

To characterise the PfAlba4-associated protein interaction network, we utilised a proximity labelling approach termed ‘BioID’, which employs a promiscuous biotinylator for tagging of proximal and interacting proteins (‘proxiome’) (Figure 3a) (Roux *et al*, 2012). Here, we utilised miniTurboID, a modified biotin ligase (BirA*), fused to a dimerisable FRB subunit to facilitate targeted, rapid and efficient biotinylation of proximal proteins (Branon *et al*, 2018a). The miniTurboID plasmid was co-transfected into the PfAlba4^2xFKBP-GFP^ transgenic line and retention of the miniTurboID episome (PfAlba4^2xFKBP-GFP/mTID-FRB-mCh^) was confirmed by western blot, followed by confirmation of rapalog-inducible dimerisation by live cell microscopy (Figure 3b, c; Supplementary figure 4a). Prior to proteomics analysis, tightly synchronised late schizont stage parasites were harvested, and biotinylation efficiency was observed by western blot (Figure 3d; Supplementary figure 4b) and immunofluorescence microscopy (Figure 3e; Supplementary figure 4c).

**Figure 3.**
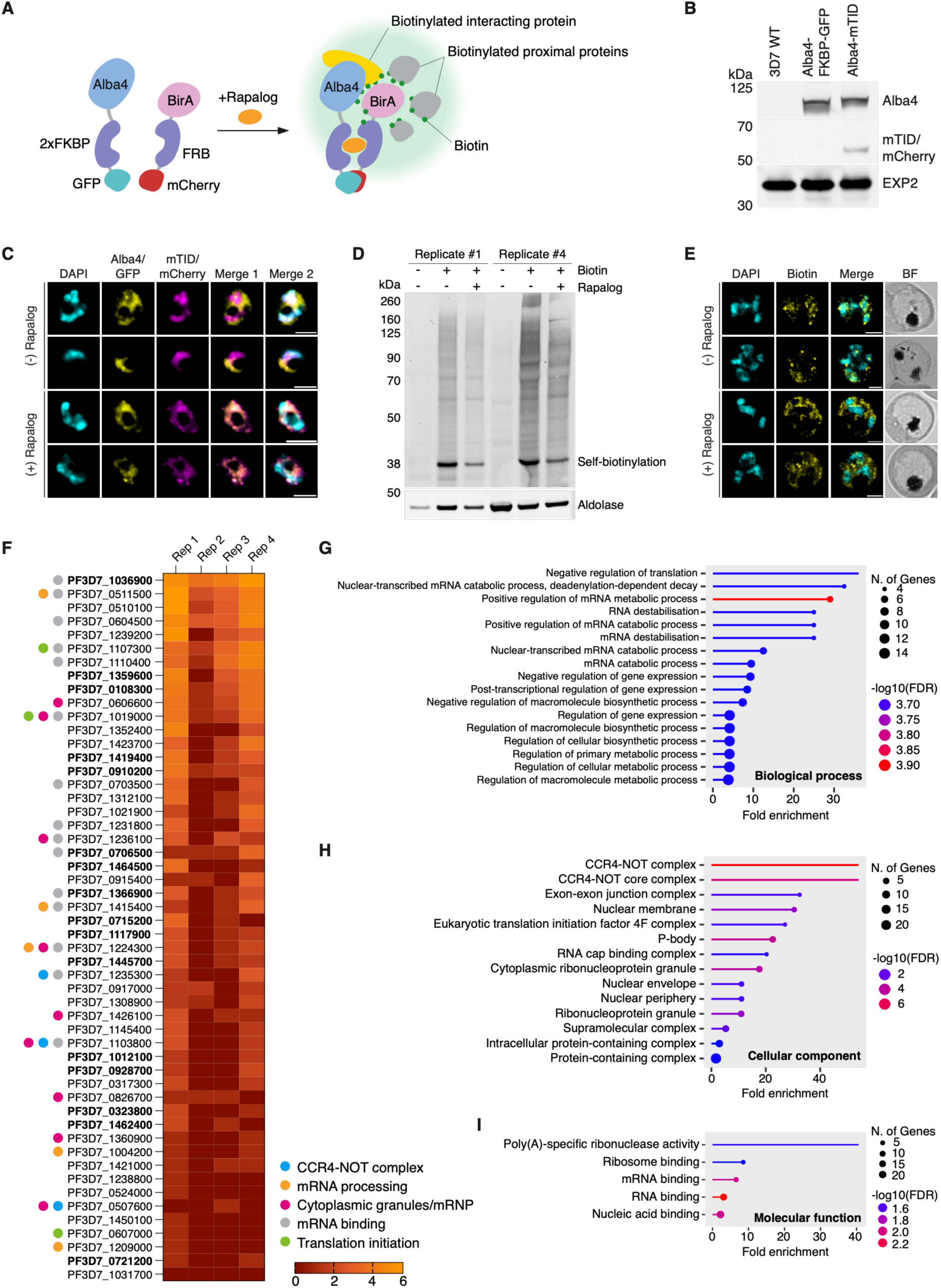
Identification of potential interacting protein partners of PfAlba4 using miniTurboID proximity labelling. **A)** Schematic representation of miniTurboID mediated proximity labelling of Alba4 proximal proteins. A miniTurboID plasmid was co-transfected into the PfAlba4^2xFKBP-GFP^ tag line (PfAlba4^2xFKBP-GFP/mTID-FRB-mCh^). The episomally expressed miniTurbo is fused to an FRB dimerisable subunit to facilitate targeted, rapalog-inducible biotinylation of proteins proximal to PfAlba4 within a 30-minute window. **B)** Western blot to confirm episome retention in the PfAlba4^2xFKBP-GFP/mTID-FRB-mCh^ line was probed with anti-GFP to detect PfAlba4, anti-mCherry to detect the miniTurbo, and anti-EXP2 was used as a loading control. **C)** Live cell imaging shows dimerisation of tagged PfAlba4 (GFP/yellow) and the mCherry tagged biotinylator (magenta). Nuclei are stained with DAPI (cyan), scale bars = 5 µm. **D)** Western blot validation of biotinylation efficiency. Samples were taken from each replicate harvest and validated by western blots probed with streptavidin for detection of biotinylated proteins. Shown are untreated (-/-), biotin only treated (+/-), and biotin and rapalog (+/+) treated samples from replicates 1 and 4. Biotinylation can be observed in the +Biotin only and +Biotin/+Rapalog lanes. Anti-aldolase (∼40 kDa) was used as a loading control. **E)** Immunofluorescence microscopy shows differential biotinylation in the presence or absence of rapalog. Methanol fixed smears from harvest replicates were stained with streptavidin (Biotin/yellow) and the nuclear stain DAPI (cyan). In the absence of rapalog, biotinylation largely overlaps with PfAlba4^2xFKBP-GFP/mTID-FRB-mCh^ line parasite nuclei, whereas in the presence of rapalog the biotin signal is detected in the parasite cytoplasm. Scale bars = 2 µm. **F)** Heat map of miniTurboID significantly enriched proteins that appeared in at least 3 replicates for the PfAlba4^2xFKBP-GFP/mTID-FRB-mCh^ line. Proteins were considered significantly enriched if they displayed a biotinylated peptide log2 fold change ≥2 and appeared in all +Biotin/+Rapalog samples or were present in two or more +Biotin/+Rapalog samples with ≥2 peptides in at least two of the replicates. Values represented are the log2 fold change of biotinylated peptides in the rapalog treated PfAlba4^2xFKBP-GFP/mTID-FRB-mCh^ samples compared to the PfAlba4^2xFKBP-GFP/mTID-FRB-mCh^ biotin only samples for each replicate. Proteins with a conserved, unknown function annotation on PlasmoDB are highlighted in bold. Additional proteins of interest include CCR4-NOT complex proteins (blue dot), proteins involved in mRNA processing (yellow dot), known and putative components of cytoplasmic granules (pink dot), mRNA binding proteins (grey dot), and proteins involved in translation initiation (green dot). **G)** Gene ontology analysis of miniTurboID high confidence proteins enriched biological processes, **H)** cellular component, and **I)** molecular function. Terms on each graph are sorted by fold enrichment. Number of genes are indicated by circle size and statistical value by colour as indicated in the associated key for each graph (ShinyGO 0.85).

Using miniTurbo BioID, we identified 67 significantly enriched potential interacting proteins for PfAlba4 (biotinylated peptide log2 fold change ≥2 and A) present in all +Biotin/+Rapalog samples or B) present in two or more +Biotin/+Rapalog samples with ≥2 biotinylated peptides in at least two of the replicates) during late blood stage development with 52 of these proteins observed in at least 3 out of 4 replicates (Supplementary figure 4d; Supplementary data 2). These proteins include known and predicted components of cytoplasmic ribonucleoprotein granules and RNA regulatory complexes and Gene Ontology enrichment analysis revealed an enrichment of proteins involved in the processing and regulation of mRNA (Figure 3f – i; Supplementary data 2).

Among the identified proteins were 6 out of the 12 annotated *P. falciparum* CCR4-NOT complex proteins, namely putative protein CAF40 (PF3D7_0507600), putative CCR4 domain-containing protein 1 (PF3D7_0519500), CCR4-associated factor 1 (PF3D7_0811300), and putative CCR4-NOT transcription complex subunits 4 (PF3D7_1235300), NOT1-G (PF3D7_1103800), and NOT1 (PF3D7_1417200). The CCR4-NOT complex is a conserved regulator of cytoplasmic mRNA turnover and repression in other organisms and is found highly conserved across *Plasmodium* spp. (Caulier *et al*, 2025; Collart, 2016). Additional candidate PfAlba4-associated proteins include putative polyadenylate-binding protein (PF3D7_1424300), polyadenylate-binding protein-interacting protein 1 (PF3D7_1107300), eukaryotic translation initiation factors subunit eIF2A (PF3D7_1019000) and EIF4G (PF3D7_1312900), and translation initiation factor IF-2 (PF3D7_0607000), further supporting an association between PfAlba4 and proteins involved in translational regulation.

Notably, many of the high confidence proteins identified for the PfAlba4 BioID lack a significant functional annotation with ∼28% having PlasmoDB annotations of conserved, unknown function (Alvarez-Jarreta *et al*, 2024). Among these, we chose two of the unannotated proteins for further investigation, PF3D7_1036900 and PF3D7_0706500.

### Functional characterisation of potential interacting partner PF3D7_1036900

PF3D7_1036900 is a 192 kDa protein with minimal annotations in PlasmoDB and a largely disordered structural prediction in AlphaFold (Supplementary figure 5a) (Jumper *et al*, 2021). An orthologue of this protein has previously been identified as a high confidence potential interacting partner of PyAlba4 (PY17X_0521000) in co-immunoprecipitation experiments, and its ortholog in *P. berghei* has been indirectly linked to PbAlba4 (PBANKA_0519900) through associations with PbDOZI and PbCITH (Mair *et al*., 2010; Muñoz *et al*., 2017).

To investigate its function, we endogenously tagged PF3D7_1036900 (PF3D7_1036900^2xFKBP-GFP^) using selection linked integration and successful integration was verified by PCR (Figure 4a). Expression of the tagged protein was then confirmed by live cell microscopy and a cytosolic localisation similar to PfAlba4 was observed (Figure 4b; Supplementary figure 5c). However, western blots probed with anti-GFP to detect the tagged protein displayed a significant amount of protein cleavage (Figure 4c). Repeated experiments with different cell lysates prepared with protease inhibitors produced the same result, indicating that this is likely endogenous protein cleavage during parasite development (Supplementary figure 5b). It was observed that the cleavage increases over the course of blood stage development, with very little full-length protein observed past trophozoite stages, indicting proteolytic cleavage increases as blood stage development progresses. Immunofluorescence assays were performed and confirmed a ubiquitous cytoplasmic localisation for this protein, similar to that of PfAlba4 (Figure 4d; Supplementary figure 5d). However, due to the levels of PF3D7_1036900 proteolytic processing observed, further localisation and attempts to identify interacting partners were not undertaken.

**Figure 4.**
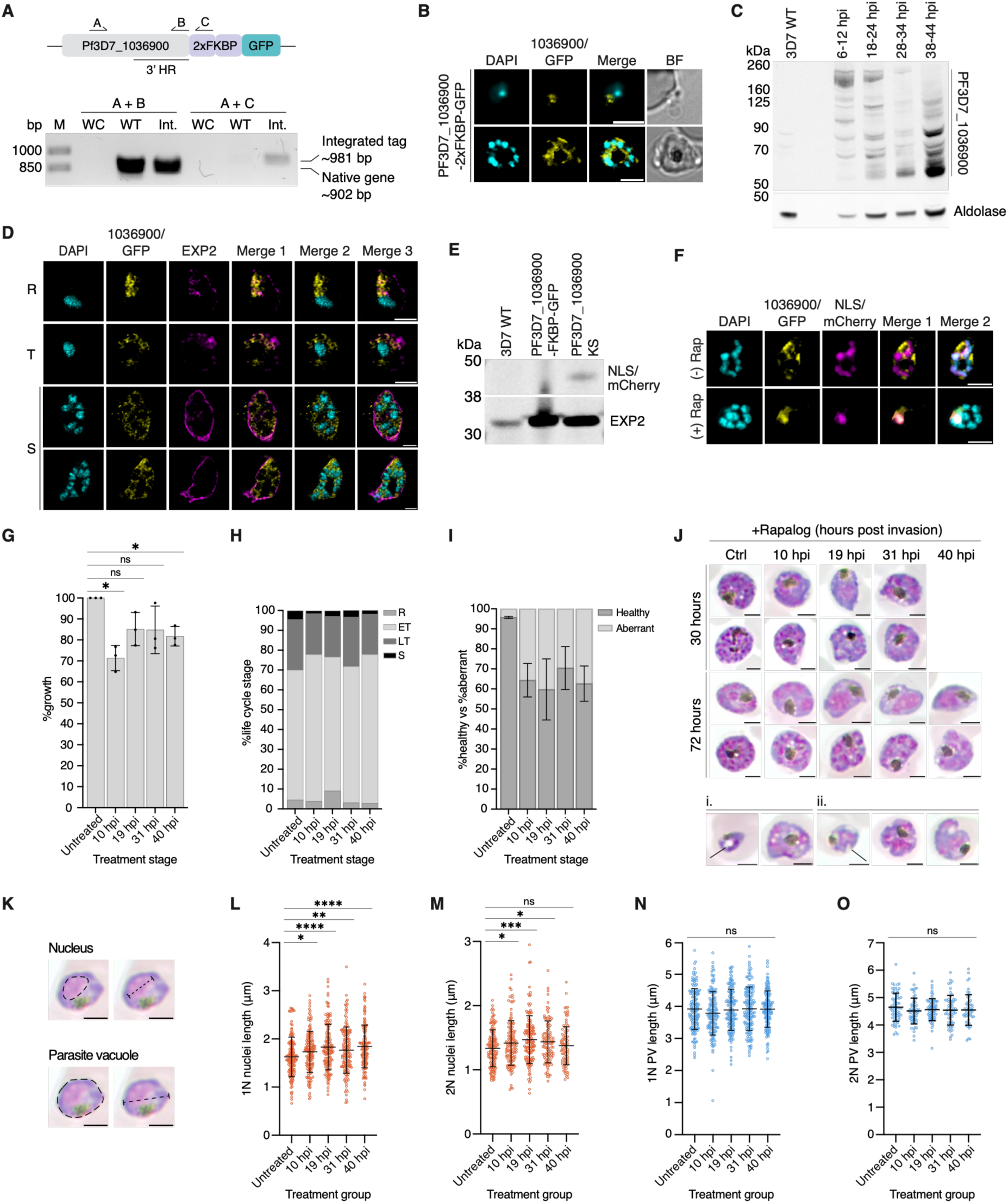
Endogenous tagging and characterisation of PF3D7_1036900. **A)** Successful endogenous tagging was confirmed by PCR for the PF3D7_1036900^2xFKBP-GFP^ line. Primers A and B correspond to the native gene sequence (902 bp) and bands of correct size were observed for the wildtype control (WT) and the integration line (Int.). Primers A and C amplify a sequence that includes the native gene and the 2xFKBP tag (981 bp), a band was only seen in the PF3D7_1036900^2xFKBP-GFP^ line, indicating successful tagging. No bands were seen in either of the water controls (WC). **B)** Live cell microscopy (Scale bars = 5 μm) and, **C)** western blot analysis confirm expression of the integrated tag in the PF3D7_1036900^2xFKBP-GFP^ line. Western blots were stained with anti-GFP to detect the tagged protein (expected size ∼244 kDa), anti-aldolase was used as a loading control and lysate from 3D7 wildtype (3D7 WT) as an additional control. PF3D7_1036900 was detected across the blood stage with progressive cleavage as the lifecycle progresses. **D)** Immunofluorescence microscopy highlights a disperse cytoplasmic localisation across blood stage development in the PF3D7_1036900^2xFKBP-GFP^ line. PF3D7_1036900 (GFP/yellow), PfEXP2 (magenta), and nuclei stained with DAPI (cyan) are shown. Merge 1 – PF3D7_1036900 and PfEXP2, Merge 2 – PF3D7_1036900 and DAPI, Merge 3 – all channels. Scale bars = 2 μm. **E)** Retention of a nuclear mislocalising episome was confirmed by western blot, creating the PF3D7_1036900^2xFKBP-GFP/NLS-FRB-mCh^ line. PF3D7_1036900 was detected by anti-GFP, mislocalising episome by anti-mCherry, and the loading control anti-EXP2 was used. **F)** Rapalog-inducible nuclear mislocalisation was confirmed by live cell microscopy (n=2) for the PF3D7_1036900^2xFKBP-GFP/NLS-FRB-mCh^ line. Merge 1 shows the PF3D7_1036900^2xFKBP-GFP^ protein in yellow (GFP) and NLS-FRB-mCherry (magenta), merge 2 shows all three channels. Nuclei were stained with DAPI (cyan) and scale bars = 5 μm. **G)** Mislocalisation of PF3D7_1036900 to the nucleus for PF3D7_1036900^2xFKBP-GFP/NLS-FRB-mCh^ rapalog treated parasites resulted in a moderate growth defect that was most pronounced when mislocalisation was initiated in ring stage parasites (n=3). Paired two-tailed t-tests were used to observe significance, ns = non-significant, \**p* <0.05. Individual data points, mean, and standard deviation are shown. **H)** Distribution of lifecycle stages in each treatment group where rapalog was added to PF3D7_1036900^2xFKBP-GFP/NLS-FRB-mCh^ parasites at ring stages (RT), early trophozoites (ET), late trophozoites (LT), or schizonts (S). **I)** Proportion of parasites deemed to be aberrant upon microscopy analysis of Giemsa smears taken at the assay end point. Healthy parasites are represented in dark grey, aberrant parasites in light grey. Error bars = standard deviation. **J)** Representative Giemsa images from the untreated PF3D7_1036900^2xFKBP-GFP/NLS-FRB-mCh^ control and rapalog treated PF3D7_1036900^2xFKBP-GFP/NLS-FRB-mCh^ parasites from each treatment time point at 30- and 72-hours post-assay set-up. Aberrant morphologies included presence of stress vacuoles in trophozoites and schizonts (i) and membrane irregularities and invagination (ii). **K)** Example measurements of nuclear material and the parasite vacuole indicated by the dotted lines. **L)** Nuclei of 1N and **M)** 2N parasites and their respective vacuoles **N)** and **O)** were measured in Fiji/ImageJ from each assay replicate end point. Measurements were taken from 50 1N parasites per treatment group/replicate. Due to their lower abundance, all 2N parasites present in Giemsa smears were measured. All rapalog treated cultures displayed significantly larger volumes of Giemsa-stained nuclear material in 1N parasites compared to the untreated control. Nucleus size was also significantly increased in three out of four treatment groups for 2N parasites. There were no statistically significant changes observed in vacuole size. P-value determined by Welch’s t-test (ns = non-significant; \**p* <0.05, \*\**p* <0.01, \*\*\**p* <0.001, \*\*\*\**p* <0.0001), error bars indicate mean and standard deviation.

PF3D7_1036900^2xFKBP-GFP^ parasites were then co-transfected with a nuclear mislocalising plasmid (PF3D7_1036900^2xFKBP-GFP/NLS-FRB-mCh^; called PF3D7_1036900-KS) and successful retention was confirmed by western blot and function confirmed by live cell microscopy in the presence or absence of rapalog (Figure 4e, f; Supplementary figure 5e). Nuclear mislocalisation assays were conducted as for PfAlba4, with mislocalisation induced at rings, early and late trophozoites, and schizont stages. Rapalog was removed at 48-hours post-assay set-up and parasite growth assessed at 72-hours. Nuclear mislocalisation of PF3D7_1036900-KS had only a modest impact on parasite growth across the blood stage compared to untreated PF3D7_1036900-KS control parasites, with the largest impact observed in ring-stage treated cultures with a ∼30% reduction in parasite growth (Figure 4g; Supplementary data 1). As with PfAlba4, no significant change in the distribution of life stages was observed compared to the untreated control (Figure 4h; Supplementary figure 6a). However, we did observe that there were obvious morphological defects in a proportion of rapalog-treated PF3D7_1036900-KS parasites compared to the untreated control, largely the presence of stress vacuoles and parasite membrane irregularities (Figure 4i, j; Supplementary figure 6b).

Notably, while assessing the morphology of the mislocalised parasites, we also observed that many of the PF3D7_1036900-KS parasites appeared to display nuclear enlargement during early trophozoite stages compared to the untreated control PF3D7_1036900-KS parasites (Supplementary figure 6b). To quantify this phenotype, the nuclei and vacuoles of 1N and 2N parasites were measured from images of Giemsa smears as demonstrated in Figure 4k. From these measurements we confirmed that there was a significant enlargement of nuclear material in single nuclei (1N) PF3D7_1036900-KS parasites across all treatment times and three out of the four treatment groups for 2N PF3D7_1036900-KS parasites compared to PF3D7_1036900-FKBP-GFP parasites (Figure 4l, m; Supplementary data 1). Interestingly, we observed no significant difference in parasite vacuole size for the 1N or 2N PF3D7_1036900-KS trophozoites (Figure 4n, o). To assess whether the increased nuclei size was not just an artefact of rapalog treatment, we also measured the nuclei of parasites for untreated and rapalog treated 3D7 wildtype parasites. In these control assays, we observed no significant increase in nuclear material, thus indicating that the phenotype in PF3D7_1036900-KS parasites is independent of the rapalog treatment (Supplementary figure 6c, d). Together, these findings implicate PF3D7_1036900 in maintaining normal parasite growth and cellular morphology, with its mislocalisation additionally associated with altered nuclear dimensions during asexual blood stage development.

### Functional characterisation of potential interacting partner PF3D7_0706500

Similar to PF3D7_1036900, PF3D7_0706500 is a large protein (196 kDa), with orthologues in *P. berghei* (PBANKA_0804100) and *P. yoelii* (PY17X_0806800) previously identified as likely proximal to Pb/PyAlba4 in proteomic studies (Mair *et al*., 2010; Muñoz *et al*., 2017). AlphaFold predicts a largely disordered protein with three small structural domains, with one of these annotated as an RNA recognition motif (RRM) domain (Supplementary figure 7a) (Jumper *et al*., 2021).

Here, we generated a PF3D7_0706500^2xFKBP-GFP^ transgenic parasite line and tag integration was validated by PCR (Figure 5a). Live cell microscopy and western blot analysis of parasite lysates from across blood stage development showed that PF3D7_0706500 is ubiquitously expressed across the blood stage, similar to PfAlba4 (Figure 5b, c; Supplementary figure 7b), and immunofluorescence assays indicated a nucleocytoplasmic localisation, with an increasingly cytoplasmic distribution during later blood stage development (Figure 5d; Supplementary figure 7c).

**Figure 5.**
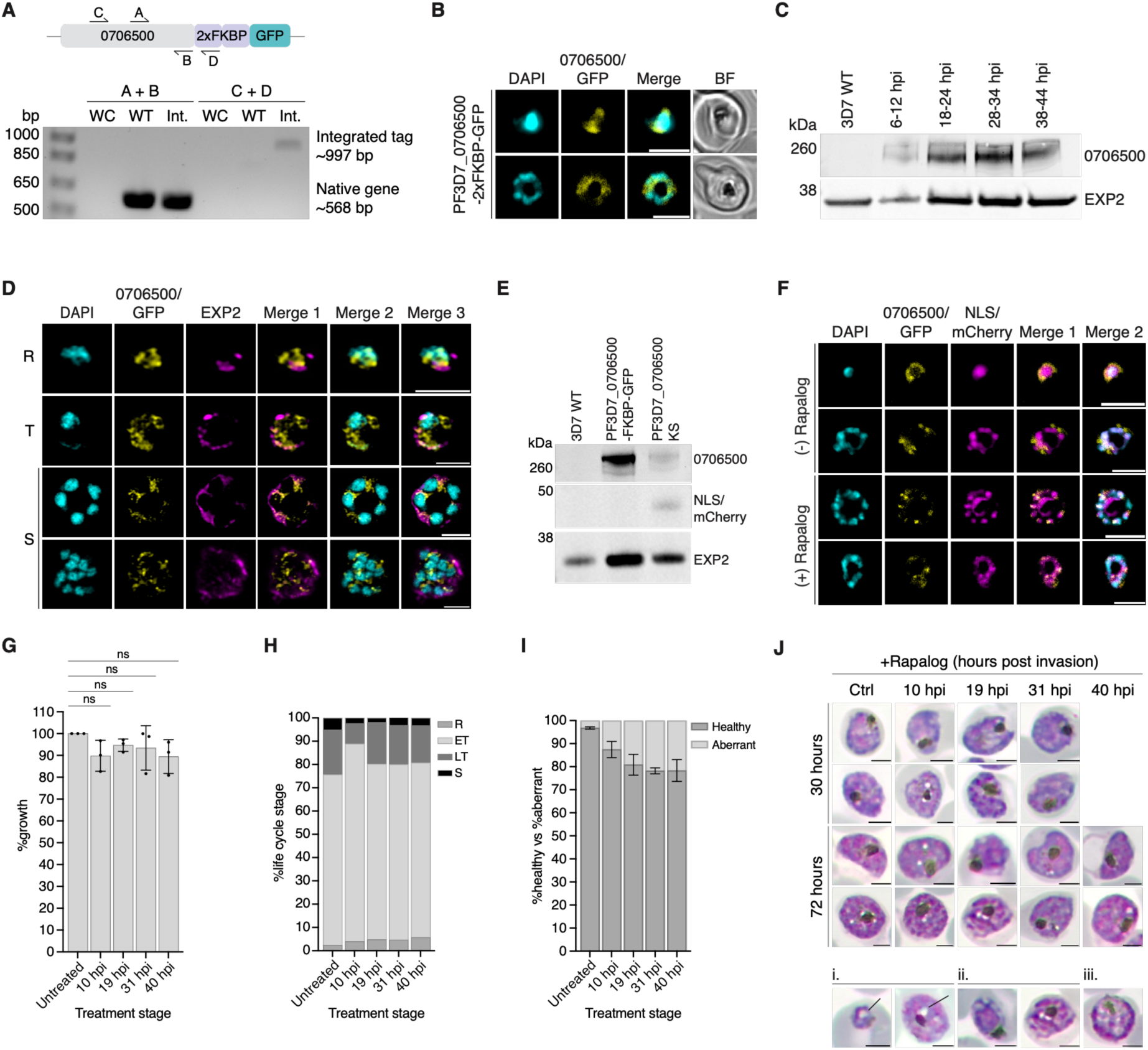
Endogenous tagging and characterisation of PF3D7_0706500. PCR confirmed successful endogenous tagging. Primers A and B amplified the native sequence (568 bp) in both the wildtype control (WT) and the PF3D7_0706500^2xFKBP-GFP^ line (Int.). Primers C and D amplified a region covering part of the native sequence and the 2xFKBP tag (997 bp). A band for the integrated tag was seen only in the integration sample. No bands were observed in either water control. **B)** Expression of the endogenous tag was confirmed by live cell microscopy. PF3D7_0706500^2xFKBP-GFP^ is shown in yellow (GFP). Nuclei were stained with DAPI (cyan), scale bars = 5 μm. **C)** Western blots further validated expression of the tagged protein and anti-GFP staining revealed a band around the expected size of ∼248 kDa for the PF3D7_0706500^2xFKBP-GFP^ line while a band was absent in the 3D7 WT control. anti-EXP2 (∼33 kDa) was used as a loading control. **D)** Immunofluorescence microscopy demonstrated ubiquitous PF3D7_0706500^2xFKBP-GFP^ expression across blood stage development with nucleocytoplasmic localisation. Parasites were stained with anti-GFP to detect the tagged protein (yellow), anti-EXP2 (parasite vacuole/magenta), and the nuclear stain DAPI (cyan). Merge 1 – PF3D7_0706500 and PfEXP2, Merge 2 – PF3D7_0706500 and DAPI, Merge 3 – all channels. Scale bars = 2 μm. **E)** Retention of a nuclear mislocalising episome was confirmed for the PF3D7_0706500^2xFKBP-GFP/NLS-FRB-mCh^ line by western blots probed with anti-GFP (PF3D7_0706500) and anti-mCherry (NLS episome) and anti-EXP2 was used as a loading control. **F)** Inducible mislocalisation was confirmed for PF3D7_0706500-KS by live cell microscopy in the presence (+) or absence (-) of rapalog. PF3D7_0706500 is represented in yellow (GFP), mislocalising episome (mCherry/magenta), nuclei are stained with DAPI (cyan), brightfield image (BF) is in grey. Scale bars = 5 μm. **G)** PF3D7_0706500 mislocalisation to the nucleus had no significant growth defects for rapalog treated PF3D7_0706500-KS parasites compared to the untreated control. Paired two-tailed t-tests, ns = non-significant. Individual data points and mean n=3 biological replicates are shown, error bars show standard deviation (SD). **H)** Quantification of parasite stages at the assay end point showed little deviation in life cycle stage across treatment groups however, **I)** a slight increase in aberrant phenotypes was observed in rapalog-treated PF3D7_0706500-KS cultures compared to the untreated PF3D7_0706500-KS parasites. Error bars = SD. **J)** Giemsa smears representative of the majority of the rapalog treated PF3D7_0706500-KS parasite population at 30-hours and 72-hours post-assay set-up. Representative images of parasites classified as ‘aberrant’ including parasites displaying obvious stress vacuoles (i), irregular membranes/morphology and cellular disorganisation (ii), and parasites with an elongated/oval-shaped morphology (iii).

We next generated a nuclear mislocalisation line for PF3D7_0706500 (PF3D7_0706500^2xFKBP-GFP/NLS-FRB-mCh^) and successful retention of the mislocalising episome by the parasite was validated by western blot with mislocalisation verified by live cell microscopy (Figure 5e, f; Supplementary figure 7d). Nuclear mislocalisation assays were set-up as for PfAlba4 and PF3D7_1036900 and parasite growth in rapalog-treated PF3D7_0706500-KS samples was compared to untreated PF3D7_0706500-KS control parasites. There was trend for reduced growth when PF3D7_0706500 was mislocalised to the nucleus, but this was not found to be significant (Figure 5g; Supplementary data 1). There was also limited differences seen with PF3D7_0706500-KS and a trend for an increasing proportion of aberrant parasites, but these measures also did not reach significance (Figure 5h, i; Supplementary figure 8a, b). Together, these findings indicate that PF3D7_0706500 mislocalisation is associated with alterations in morphology for some parasites leading to limited impacts on blood stage growth, suggesting PF3D7_0706500s functional contribution is somewhat distinct from that observed for PfAlba4.

### Coimmunoprecipitation and identification of PF3D7_0706500 proximal proteins

To further investigate the potential association between PF3D7_0706500 and PfAlba4, we next utilised the GFP tag of PF3D7_0706500-FKBP-GFP to undertake coimmunoprecipitation (CoIP) experiments followed by proteomics analysis to identify potential interacting partners of PF3D7_0706500 (Figure 6a). This approach identified 161 significantly enriched (log2 fold change ≥2 and adjusted *p*-value ≤0.05) potential interacting partners for PF3D7_0706500, indicating a rich network of proximal proteins (Figure 6b; Supplementary data 3). PfAlba4 was enriched in the coimmunoprecipitation data with high confidence (1.37 log2 fold change, 0.003 adjusted *p*-value), but did not meet the log2 fold change cut-off to be considered significant. Ten enriched proteins were identified to directly overlap between the PfAlba4 miniTurboID data and PF3D7_0706500 CoIP datasets (Figure 6c). Notably, 3 out of the 6 PfCCR4-NOT complex subunits, namely CAF1, NOT1, and NOT1-G, identified in the PfAlba4 miniTurboID data were also highly enriched in the PF3D7_0706500 CoIP data, alongside the putative PABP1. Other shared proteins included putative KH domain- and WD repeat-containing proteins (PF3D7_0510100 and PF3D7_0606600 respectively), both with predicted roles in RNA/mRNA binding and/or processing, as well as asparagine-rich antigen Pfa35-2 (PF3D7_0105700), and two conserved proteins of unknown function (PF3D7_0323800 and PF3D7_1012100).

**Figure 6.**
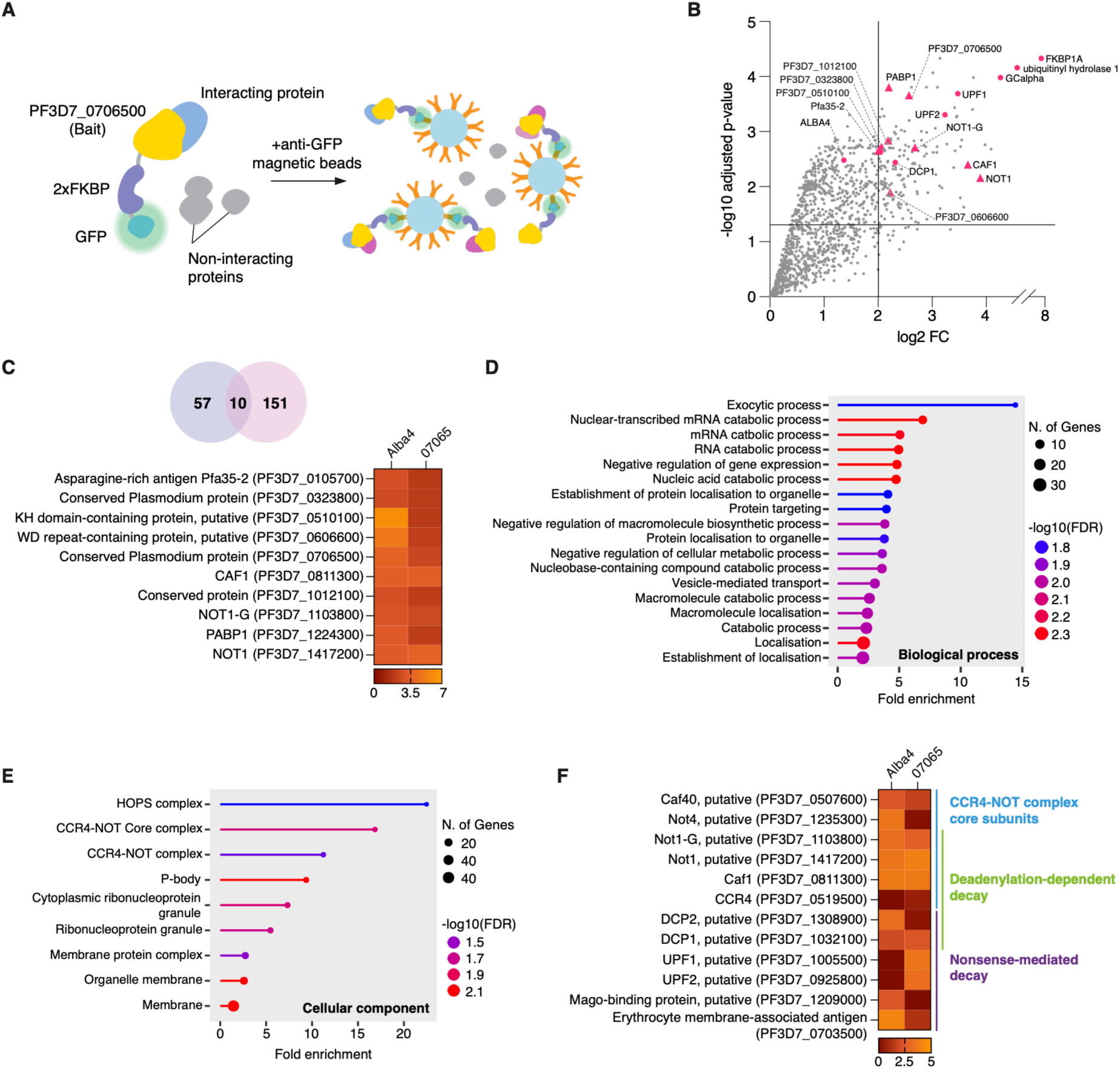
Identification of PF3D7_0706500 potential interacting partners by coimmunoprecipitation. **A)** Schematic representation of coimmunoprecipitation (CoIP). The anti-GFP conjugated magnetic beads bind to the GFP-tagged protein of interest (PF3D7_0706500) in PF3D7_0706500^2xFKBP-GFP^ parasites to pulldown this protein as well as any stably interacting protein partners which are then analysed by mass spectrometric analysis. **B)** Volcano plot showing the proteins identified in PF3D7_0706500 CoIP experiments (n=4 biological replicates). The -log10 transformed adjusted p-values are represented on the y-axis and log2 fold-change (log2 FC) on the x-axis. Proteins with a log2 FC of ≥2 for PF3D7_0706500^2xFKBP-GFP^ parasites (compared to 3D7 wildtype control) and a p-value of ≤0.05 were considered statistically significant (161 proteins). Pink triangles indicate proteins overlapping with the Alba4-miniTurboID significant proteins. Pink circles indicate additional enriched proteins including the endogenous tag component FKBP1A and the two highest enriched proteins ubiquitinyl hydrolase 1 (PF3D7_1408700) and guanylyl cyclase alpha (GCalpha, PF3D7_1138400) and additional proteins involved in negative regulatory processes including putative regulators of nonsense transcripts UPF1 and UPF2 (PF3D7_0925800, PF3D7_1005500) and putative mRNA-decapping enzyme DCP1 (PF3D7_1032100). **C)** 10 proteins were observed to overlap between the high confidence Alba4-miniTurboID and high confidence PF3D7_0706500 CoIP data. The heatmap represents the log2 FC for these proteins in each experiment. **D)** Enriched biological processes and, **E)** Cellular components identified by Gene Ontology enrichment analysis (ShinyGO 0.85) for the high confidence PF3D7_0706500^2xFKBP-GFP^ CoIP data. Terms are sorted by foldchange, FDR is indicated by colour, and protein number by circle size as indicated in their respective legends. The HOPS complex was the highest enriched cellular component and is known to be involved in endocytic and trafficking processes (Mesén-Ramírez *et al*, 2025). **F)** Heatmap highlighting some proteins involved in negative regulation of translation that were enriched in either one or both of the proteomics datasets. Highlighted are proteins with known or predicted involvement in CCR4-NOT complex, deadenylation-dependent decay, and nonsense-mediated decay. Values represented are the log2 FC for each protein in the PfAlba4 and PF3D7_0706500 datasets.

Gene ontology analysis revealed shared enrichments of CCR4-NOT complex, P-body, and more broadly, cytoplasmic ribonucleoprotein granule proteins in the PF3D7_0706500 CoIP and PfAlba4 miniTurboID datasets (Figure 6d, e; Supplementary data 3). Collectively, these datasets identified six PfCCR4-NOT complex subunits, alongside putative mRNA-decapping enzyme subunits DCP1 (PF3D7_1032100) and DCP2 (PF3D7_1308900) as well as putative regulator of nonsense transcript proteins UPF1 (PF3D7_1005500) and UPF2 (PF3D7_0925800), all proteins with putative roles in deadenylation-dependent decay and nonsense-mediated decay (Figure 6f).

The PF3D7_0706500 CoIP data also revealed a distinct enrichment of homotypic fusion and vacuole protein sorting (HOPS) complex, a part of the larger HOPS/CORVET (class C core vacuole/endosome tethering) complex that performs roles in protein transport. Studies in *P. falciparum* have demonstrated both canonical and parasite-specific roles for this machinery during blood stage development (Mesén-Ramírez *et al*, 2025).

Together, these datasets revealed overlapping associations with RNA regulatory proteins and pathways, alongside several conserved proteins of unknown function that may represent uncharacterised components of these regulatory networks. The distinct enrichments further suggest that PfAlba4 and PF3D7_0706500 may participate in both shared RNA-regulatory processes and separate cellular pathways.

### RIP-seq identifies PfAlba4-associated transcripts in late blood stage development and demonstrates a preference for pyrimidine-rich RNA elements

Given that PfAlba4 is annotated as a DNA/RNA-binding protein and its association with RNA regulatory proteins in proximity ligation assays, we next wanted to identify potential mRNA substrates to further understand the regulatory roles of these proteins. RIP-seq identified 692 significantly enriched (≥2.5-fold enrichment and FDR ≤0.05) PfAlba4-associated transcripts from 5,043 transcripts evaluated in schizont-stage parasites (Figure 7a; Supplementary data 4), with representative transcripts reflecting varying degrees of enrichment following PfAlba4 immunoprecipitation (Figure 7b). To identify putative cis-regulatory elements mediating PfAlba4 target recognition, we performed discriminative motif analysis (XSTREME) on the 3′UTRs of PfAlba4 RIP-seq targets against non-target *P. falciparum* 3′UTRs as background. MEME identified a pyrimidine-rich element (CYYYCMUC; Figure 7c) enriched 1.69-fold in target sequences relative to background (6.76% vs 4.06%; SEA P = 4.62 × 10⁻³, E-value = 4.62 × 10⁻², q = 0.023) (Figure 7c). The motif is dominated by pyrimidine residues (Y = C or U), consistent with the recently described preference of ALBA4 proteins in *Arabidopsis* for uridine- and pyrimidine-rich RNA elements (Reichel *et al*, 2024). Tomtom comparison against the Ray2013 RNAcompete database identified significant similarity to the binding profile of PCBP2 (E = 9.9 × 10⁻⁸), which recognises C/U-rich sequence elements (Supplementary data 5) (Ray *et al*, 2013). Together, these findings identify a pyrimidine-rich 3′UTR element enriched within a subset of PfAlba4-associated transcripts, consistent with sequence-dependent contributions to PfAlba4 target recognition.

**Figure 7.**
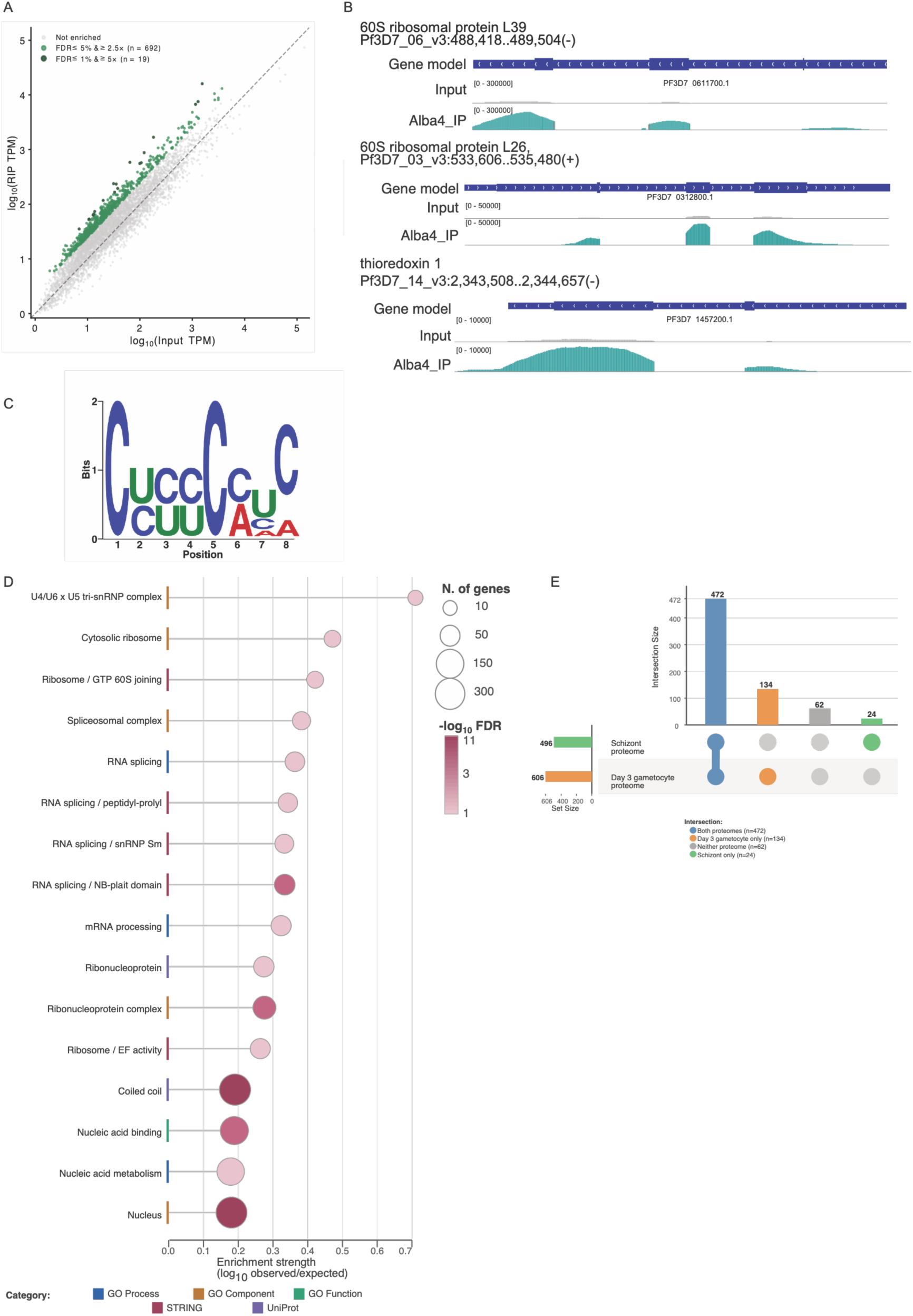
RIP-Seq identifies 692 PfAlba4-associated mRNA targets in *P. falciparum* schizonts. **A)** Scatter plot comparing log₁₀-transformed TPM values between the Alba4 RIP pulldown (RIP TPM, y-axis) and total RNA input (Input TPM, x-axis) for all genes detected in the RIP-Seq experiment. Grey points represent non-enriched transcripts. Green points represent primary targets enriched at FDR ≤ 5% and ≥ 2.5-fold (n = 692), with dark green points indicating high-confidence targets at FDR ≤ 1% and ≥ 5-fold enrichment (n = 19). The dashed line indicates equal abundance between pulldown and input. **B)** Representative IGV genome browser tracks showing PfAlba4 RIP-Seq enrichment over three example target genes. Tracks display the gene model (top), total RNA input (middle, grey), and Alba4 immunoprecipitate (bottom, teal) for 60S ribosomal protein L39 (PF3D7_0611700; rank 1, 10.4×), 60S ribosomal protein L26 (PF3D7_0312800; rank 2, 9.3×), and thioredoxin 1 (PF3D7_1457200; rank 3, 9.1×). Read depth scales are indicated in brackets. **C)** Sequence logo of the top-ranked MEME-7 motif (CYYYC MUC) identified by discriminative XSTREME analysis of PfAlba4 target 3’UTRs against 3,335 non-target *P. falciparum* 3’UTRs as background (MEME Suite v5.5). Letter height represents information content in bits (maximum 2 bits). The motif is a pyrimidine-rich element enriched 1.69-fold in target sequences relative to background (6.76% vs 4.06%; SEA p = 4.62 × 10⁻³, E-value = 4.62 × 10⁻²) and shows significant Tomtom similarity to the binding profile of PCBP2 (poly-C binding protein 2; E = 9.9 × 10⁻⁸), consistent with the recently described preference of Alba4 proteins for pyrimidine-rich elements in 3’UTRs. **D)** Horizontal lollipop plot showing functional enrichment of PfAlba4 RIP-Seq targets. The x-axis shows enrichment strength (log₁₀ observed/expected gene count). Dot size is proportional to the number of observed genes and dot colour indicates −log₁₀ FDR (light pink: FDR ∼0.04; dark red: FDR < 0.001). Categories are colour-coded as indicated. Enrichment analysis was performed using STRING (v11.5) against a background of all *P. falciparum* 3D7 genes. Notable enriched categories include spliceosomal components (U4/U6×U5 tri-snRNP complex, spliceosomal complex, RNA splicing), ribosome biogenesis, and ribonucleoprotein complexes. **E)** UpSet plot comparing PfAlba4 RIP-Seq targets (n = 692) against stage-specific proteomes from *P. falciparum* schizonts and wild-type Day 3 gametocytes. The upper bar chart shows intersection sizes; the dot matrix below indicates which proteome sets are active in each intersection. Horizontal bars on the left show total set sizes. 472 targets (68.2%) were detected at the protein level in both stages, 134 (19.4%) exclusively in Day 3 gametocytes, 24 (3.5%) exclusively in schizonts, and 62 (9.0%) were undetected in either proteome. Birnbaum J, Flemming S, Reichard N, Soares AB, Mesén-Ramírez P, Jonscher E, Bergmann B, Spielmann T, Birnbaum J, Flemming S *et al* (2017) A genetic system to study Plasmodium falciparum protein function. *Nature Methods 2017 14:4* 14 Mesén-Ramírez JP, Fuchs G, Burmester J, Farias GB, Alape-Flores AM, Singla S, Alder A, Cubillán-Marín J, Castro-Peña C, Lemcke S *et al* (2025) HOPS/CORVET tethering complexes are critical for endocytosis and protein trafficking to invasion related organelles in malaria parasites. *PLoS pathogens* 21: e1013053

Using an FDR <0.05 threshold, functional enrichment analysis showed that PfAlba4-associated transcripts were enriched for RNA splicing, mRNA processing, nucleic acid metabolism and nucleic acid binding, with strong enrichment of nuclear and ribonucleoprotein-complex components (Figure 7d; Supplementary data 6). Enriched cellular components included the spliceosomal complex, cytosolic ribosome and U4/U6·U5 tri-snRNP, while network analysis further highlighted RNA-splicing proteins, snRNP components and translation-associated factors (Figure 7d; Supplementary data 6). Together, these analyses identify RNA processing, splicing and ribonucleoprotein function as prominent features of the PfAlba4 target set. Significant PfAlba4 targets included ribosomal proteins such as L39 (PF3D7_0611700) and L26 (PF3D7_0312800), pre-mRNA-splicing factors, RNA-binding proteins and the spliceosomal protein SNU23 (PF3D7_1243100). Other notable targets included ApiAP2 regulators such as AP2-O (PF3D7_1143100), AP2-O2 (PF3D7_0516800), AP2-P (PF3D7_1107800) and AP2-LT (PF3D7_0802100), together with chromatin-associated factors including ISWI chromatin-remodelling complex ATPase (PF3D7_062460), heterochromatin protein 1 (PF3D7_1220900), PHD finger protein, PHD2 (PF3D7_1433400) and histone H2A.Z (PF3D7_0320900). These findings indicate that PfAlba4-associated transcripts are enriched for genes encoding components of RNA-processing machinery and transcriptional and chromatin-associated regulatory pathways.

Comparison with schizont and day 3 gametocyte-specific proteomes showed that 496 of the 692 PfAlba4-associated transcripts (71.7%) had corresponding protein evidence in schizonts, leaving 196 without detectable protein products at this stage (Figure 7e; Supplementary data 7) (Chisholm *et al*, 2026; Marapana *et al*, 2026). Of these 196 transcripts, 134 (68.4%) were detected at the protein level in wild-type Day 3 gametocytes. Consequently, 630 of 692 PfAlba4 targets (91.0%) had protein-level evidence across the schizont and Day 3 gametocyte datasets, while 62 remained undetected in either proteome (Figure 7e; Supplementary data 7). These comparisons demonstrate substantial stage-dependent variation in the protein detection of PfAlba4-associated transcripts and show that most targets absent from the schizont proteome are detectable during early gametocyte development.

## Discussion

Endogenous tagging of *Plasmodium* Alba proteins has proved challenging to date and to our knowledge, the only successful instance of this has been the GFP-tagging of *P. yoelii* Alba4. Here, we successfully introduced a C-terminal endogenous tag to PfAlba4, generating a transgenic parasite line that enabled additional functional characterisation of this protein, and its associated protein and RNA interaction networks during *P. falciparum* asexual blood stage development.

In this study we observed a ubiquitous expression and cytoplasmic localisation for PfAlba4 throughout blood stage development, consistent with previous work which utilised in-house antibodies (Chêne *et al*., 2012; Min *et al*, 2024) or episomal overexpression systems (Acharya *et al*., 2025). Although PfAlba4 endogenous tagging was successful, repeated attempts to truncate the C-terminal Maf1-like domain of PfAlba4 did not yield viable parasites, indicating that this domain may be important for parasite viability. Furthermore, conditional nuclear mislocalisation of PfAlba4 resulted in reduced parasite growth and pronounced morphological abnormalities, including stress vacuoles and membrane irregularities, without significant changes in the developmental-stage distribution. These findings support an important role for PfAlba4 in maintaining normal parasite growth and cellular integrity although the molecular basis of these defects remain unresolved. Interestingly, a previous study in this parasite observed that overexpression of PfAlba4 lead to an increase in parasite growth (Acharya *et al*., 2025), further supporting the functional importance of this protein during asexual blood stage development. Nevertheless, unsuccessful domain truncation and conditional mislocalisation does not conclusively establish essentiality. Generation of a PfAlba4 inducible knock-out line would be a reasonable next step to further clarify this protein’s importance to parasite blood stage development.

Proteomics analysis revealed an extensive network of proximal proteins for PfAlba4, including the uncharacterised protein PF3D7_0706500. Subsequent CoIP analysis of PF3D7_0706500 identified several proteins shared with the PfAlba4 miniTurboID dataset, although PfAlba4 itself was only moderately enriched and didn’t meet the predefined foldchange threshold (log2 FC ≥2, adj. *p*-value ≤ 0.05). Notably, several uncharacterised proteins within these networks have predicted roles in mRNA processing and surveillance. Among these, PF3D7_0606600 (putative WD repeat-containing protein) was enriched in both datasets, and based on PlasmoDB predicted functions, is a potential p-body constituent and may have functional roles in mRNA decapping and degradation (Alvarez-Jarreta *et al*., 2024). Another candidate, PF3D7_1209000, contains a Partner of Y14 and Mago (PYM) domain, which is associated with the exon junction complex (EJC) in other organisms and is implicated in mRNA transport, nonsense-mediated decay, and translation (Bono *et al*, 2004; Gehring *et al*, 2009). Although repeated attempts to endogenously tag PF3D7_1209000 in this study were unsuccessful (data not shown), its identification further supports the association of PfAlba4 with proteins potentially involved in post-transcriptional regulation.

It is notable that orthologues of both PF3D7_0706500 and PF3D7_1036900, identified as high-confidence potential interacting partners in the PfAlba4 miniTurboID data, have been identified as either directly or indirectly associated with the *P. berghei* and *P. yoelii* orthologs of PfAlba4 (Mair *et al*., 2010; Muñoz *et al*., 2017). These findings suggest that Alba4 associated protein networks may be conserved across *Plasmodium* species, reflecting conserved cellular functions. On this basis, we undertook further characterisation of these two proteins.

Endogenous tagging of PF3D7_0706500 revealed a nucleocytoplasmic localisation throughout blood stage development, with a predominantly cytoplasmic distribution in schizont stage parasites, consistent with its identification in the PfAlba4 proximal proteome. Unlike PfAlba4, conditional PF3D7_0706500 mislocalisation did not significantly affect parasite growth, although morphological defects such as stress vacuoles and membrane irregularities were observed that bared similarity to PfAlba4-KS parasites. These findings suggest that PF3D7_0706500 may contribute to cellular regulation without being required for normal parasite proliferation under the condition’s examined.

The PF3D7_0706500 CoIP and PfAlba4 miniTurboID datasets revealed overlapping enrichment of CCR4-NOT complex components with six of the 12 *P. falciparum* CCR4-NOT complex proteins identified in both datasets. Notably, PfCAF1, PfNOT1, and PfNOT1-G were enriched in both datasets. PfCAF1 has previously been identified as an important regulator of asexual parasite egress and invasion genes (Balu *et al*, 2011). Furthermore, studies in *P. yoelii* have demonstrated distinct developmental requirements for the NOT1 paralogues, with PyNOT1 required for asexual blood stage development, while PyNOT1-G appears essential for gametocyte maturation and zygote development in this rodent malaria species (Hart *et al*, 2021). These findings suggest that the association of PfAlba4 and PF3D7_0706500 with CCR4-NOT components may be relevant to post-transcriptional regulation during parasite development.

Homologues of the canonical nonsense-mediated decay (NMD) factors UPF1 and UPF2 were also identified in the PF3D7_0706500 CoIP dataset. Although, PfUPF1 and PfUPF2 have been reported to be dispensable for *P. falciparum* asexual blood stage growth, their functional role remains poorly understood (McHugh *et al*, 2023). Notably, McHugh *et al*. (2023) also identified PfAlba4 and PF3D7_0703500 as high confidence interactors of PfUPF1/2, providing independent evidence of their association with proteins implicated in RNA surveillance. PF3D7_0703500, which was also identified in both proteomic datasets in the present study, is annotated as an erythrocyte membrane-associated antigen despite predicted nucleic acid-binding and 5’-3’ DNA helicase activity, thus the evidence generated in this and other studies suggests that it is unlikely to be associated with the erythrocyte membrane. Together, these findings suggest that the PfAlba4-associated protein network encompasses components implicated in multiple aspects of mRNA turnover and surveillance. In addition to the shared enrichment of RNA-regulatory proteins, the PF3D7_0706500 CoIP data revealed enrichment of proteins associated with the homotypic fusion and protein sorting (HOPS) complex and vesicle-mediated transport. These findings raise the possibility that PF3D7_0706500 may also associate with intracellular trafficking pathways in the parasite cytosol, though its functional involvement in these processes remains to be established.

The function of PF3D7_1036900 and its potential interactions with PfAlba4 remains unresolved. Previous studies in *P. falciparum* have linked PF3D7_1036900 to a divergent RNA N6-methyladenosine methyltransferase complex involved in the post-transcriptional regulation of mRNA (Baumgarten *et al*, 2019). In this study the authors also identified karyopherin beta (PF3D7_0524000) which was a high confidence protein in our data also. Similar to orthologues in other organisms, the *Plasmodium* karyopherin beta is a protein involved in nuclear import pathways (Mohmmed *et al*, 2003). Gene Ontology annotations in PlasmoDB for PF3D7_1036900 are limited, but propose a possible nuclear localisation and mRNA binding function. Given the endogenous protein cleavage indicated in our data, one potential hypothesis is that multiple isoforms of this protein are produced via proteolytic cleavage during parasite development and these may be localising to different subcellular compartments or have differing functions. However, it may also be that the cleavage pattern we observed is the result of normal protein turnover in the parasite and further investigation of PF3D7_1036900 and its function would need to be undertaken to resolve this.

Interestingly, despite the enrichment of PfAlba4 in the proximity labelling experiments, we did not detect any biotinylated peptides for PfAlba4 itself. However, there are several viable hypotheses to explain this. Analysis of the individual peptide data for these experiments showed that there was a significant enrichment of PfAlba4 non-biotinylated peptides in the +Biotin/+Rapalog samples compared to the +Biotin only controls (log2 fold-change of 3.91, p-value of 0.01). There were 16 peptide sequences identified and 13 of these were unique in the +Biotin/+Rapalog samples only while the remaining three sequences were also present in the +Biotin only control. Several of these sequences also overlapped, indicating specific enrichment in some regions and when mapped to the whole protein sequence for PfAlba4, amino acids 226-240 were the highest enriched residues (Supplementary figure 9a). Given this, it is plausible that PfAlba4 was largely inaccessible to the biotinylating enzyme but that non-biotinylated peptides were pulled down during enrichment due to stable interactions with biotinylated proteins. Further supporting this hypothesis, when we mapped the PfAlba4 peptides identified in the PF3D7_0706500 coimmunoprecipitation experiments in the same manner, we found that these same residues were enriched here also (Supplementary figure 9b).

Previous studies investigating the relationship between *Plasmodium* Alba4 and mRNA regulation have primarily examined changes in transcript abundance following Alba4 overexpression or genetic disruption (Acharya *et al*., 2025; Muñoz *et al*., 2017). Here, we utilised a RIP-seq approach using endogenously tagged PfAlba4 as bait to directly identify 692 significantly enriched PfAlba4-associated transcripts in late blood stage development. Subsequent enrichment analysis revealed functional enrichments for RNA processing, pre-mRNA splicing, ribonucleoprotein complexes, and translation. Notably, these included transcripts encoding ApiAP2 transcription factors and chromatin-associated regulators, suggesting that PfAlba4 may contribute to the post-transcriptional regulation of proteins involved in broader developmental gene-expression programmes. Furthermore, the identification of a pyrimidine-rich 3′UTR motif (CYYYCMUC; Y = C or U, M = A or C) is particularly interesting given recent findings in *Arabidopsis thaliana*, where ALBA4 associates with uridine- and pyrimidine-rich RNA elements and facilitates YTHDF-mediated recognition of m6A-modified transcripts (Reichel *et al*., 2024). Although this motif was enriched in only a small proportion of PfAlba4-associated transcripts, the similarity suggests that recognition of pyrimidine-rich RNA elements may represent a conserved feature of ALBA-mediated RNA regulation. Comparison with stage-specific proteomes revealed that 134 PfAlba4-associated transcripts lacking detectable protein products in schizonts had corresponding protein evidence in Day 3 gametocytes. While this observation may reflect developmental differences in protein expression or detection, it raises the possibility of stage-dependent regulation of transcript fate, consistent with the importance of mRNA storage and translational control during *Plasmodium* development (Mair *et al*., 2010; Muñoz *et al*., 2017).

Particularly noteworthy was the enrichment of transcripts encoding three CCR4-NOT components, PfNOT1, CCR4-2 (PF3D7_1363500) and CCR4-4 (PF3D7_0319200), with PfNOT1 also identified in both the PfAlba4 miniTurboID and PF3D7_0706500 CoIP datasets. This convergence of transcriptomic and proteomic evidence raises the possibility that PfAlba4-associated ribonucleoprotein complexes are linked to CCR4–NOT-mediated regulation of mRNA stability and translation, potentially including the regulation of transcripts encoding components of this machinery. Although the functional consequences of these associations remain to be established, the combined findings support a model in which PfAlba4 participates in post-transcriptional regulatory networks that may influence mRNA fate and developmental gene expression in *P. falciparum*.

Taken together, these data provide a detailed overview of PfAlba4 cellular interactions during late blood stage development and establish PfAlba4 in multiple important regulatory pathways including RNA processing and degradation. We further suggest that PfAlba4 may function as a central scaffolding protein or adaptor facilitating the docking of effector proteins mediating essential regulatory processes such as mRNA degradation.

## Methods

### Continuous culture of *P. falciparum* parasites

*P. falciparum* 3D7 wildtype parasites and all transgenic parasite lines were maintained as previously described (Trager & Jensen, 1976). Parasites were grown at 3% haematocrit in O+ human red blood cells (Lifeblood, Australia) and RPMI-HEPES supplemented with 367 µM hypoxanthine, 25 µg/mL gentamicin, 0.17% NaHCO_3_, and 0.5% Albumax II at a pH of 7.4. Cultures were stored in sealed boxes under atmospheric conditions of 1% O_2_, 5% CO_2_, and 94% N_2_ and incubated at 37^०^C.

For proximity labelling assays, parasites were cultured in biotin-free RPMI media supplemented with 1% w/v sodium bicarbonate, 0.5% Albumax II, 25 µg/mL gentamicin, 1M HEPES, and 367 µM hypoxanthine for at least two rupture cycles under standard gas and incubation conditions.

### Construction of transfection plasmids for endogenous tagging

Details of all primers used in this study can be found in Supplementary table 1.

Plasmids for endogenous tagging of target genes were constructed as previously described (Birnbaum et al., 2017). Primers were used to insert restriction sites and PCR amplify C-terminal homology regions of PF3D7_1347500, PF3D7_0706500 or PF3D7_1036900 from 3D7 wildtype gDNA.

Amplified inserts and the plasmid vector, pSLI-2xFKBP-GFP, were digested with NotI and AvrII (NEB) for endogenous tagging or NotI and MluI (NEB) for the targeted gene deletion plasmid followed by ligation. Plasmids were then screened by PCR and sequencing prior to transfection to ensure successful ligation.

### Transfection of *P. falciparum* parasites

Parasites were transfected using the schizont-loading method (Moon et al, 2013). Briefly, parasite cultures were synchronised with 150 nM of the egress inhibitor ML10 and late stage schizonts purified by 70% Percoll centrifugation (Rivadeneira et al, 1983).

For each transfection, 100 µg of plasmid DNA was ethanol precipitated then resolubilised in 10 μl of 1X TE buffer and added to schizonts with 80 µl of cytomix (0.895% KCl, 0.0017% CaCl_2_, 0.076% EGTA, 0.102% MgCl_2_, 0.0871% K_2_HPO_4_, 0.068% KH_2_PO_4_, 0.708% HEPES). Transfection mixture was transferred to a 0.2 cm BioRad cuvette and electroporated at 0.31 kV/capacitance of 960 µF. Transfected parasites were then transferred to 5 mL dishes and incubated O/N.

For retention of selection-linked integration or targeted-gene deletion episomes, transfected ring stage parasites were treated with 5 nM WR99210 (Jacobus Pharmaceuticals). For integration of the endogenous tag once stable episome retention had been achieved, 400 µg/mL G418 (Thermofisher) was added to encourage tag integration. All integration lines were assessed by PCR of genomic DNA to confirm successful integration (Supplementary Table 4).

Retention of episomal expression plasmids was selected for by 5 µg/mL blasticidin-S-deaminase HCl (Life Technologies) for the miniTurbo plasmid (miniTurbo-FRB-mCherry-BSD1^SF3A2^) and 0.9 µM of DSM1 (BEI resources) for nuclear mislocalisation plasmids (1xNLS-FRB-mCherry-DHODH^NMD3^) (Birnbaum et al., 2017; Branon et al, 2018b).

### Saponin lysis of parasite cultures

Parasite cultures were harvested by 0.15% (w/v) saponin (ThermoFisher Scientific) lysis with 1X protease inhibitor (Cømplete, Roche) for 10 min on ice. Lysed samples were then centrifuged at 16,000 x g for 10 min at 4°C and washed once in 0.075% (w/v) saponin/1X protease inhibitor. Lysates were then washed 3 times in 1X PBS/1X protease inhibitor and frozen at -80°C.

### SDS-PAGE and Western blot

Parasite lysates were DNase treated (Qiagen) and resuspended in reducing sample buffer (0.125 M Tris-HCl pH 7, 20% (v/v) glycerol, 4% (v/v) SDS, 10% (v/v) β-mercaptoethanol (Sigma-Aldrich), 0.002% (w/v) bromophenol blue (Sigma-Aldrich)). Reduced samples were then separated on 4-12% Bis-Tris pre-cast gels (Invitrogen) in 1X MES running buffer (Bolt) at 110 V/90 min. Protein was transferred to nitrocellulose membranes (BioRad, Invitrogen) for 7-10 min at 25 V. Membranes were blocked with 3% skim milk/1X PBS for 1-hour RT with gentle agitation. Blocking buffer was replaced by primary antibodies prepared in antibody diluent (0.5% skim milk/1X PBS-T 0.05%) to the following concentrations and incubated O/N at 4°C: 1:7000 mouse or rabbit anti-EXP2 (Bullen *et al*, 2012), 1:1000 mouse anti-GFP (Sigma-Roche/Invitrogen), 1:5000 rabbit anti-aldolase (Invitrogen), 1:2000 mouse anti-FKBP (Santa Cruz), 1:2000 anti-streptavidin IR800CW (LI-COR Biosciences), 1:2000 rabbit anti-mCherry (BioVision).

The following day, membranes were washed three times in wash buffer (1x PBS-T 0.05%) and secondary antibodies were added at the following concentrations in antibody diluent: 1:10,000 goat anti-mouse IR800CW (LI-COR Biosciences) and 1:15,000 goat anti-rabbit IR680RD (LI-COR Biosciences) and incubated in the dark for 1-hour RT. Membranes were washed two more times in wash buffer and once in 1X PBS, then imaged using a BioRad ChemiDoc imager. All image adjustments and volume quantifications were undertaken using the BioRad ImageLab software (version 6.1). For volume quantification, local background subtraction was used.

### Mislocalisation assays

Parasite mislocalisation lines were treated with 150 nM of ML10 at ∼ 32 h.p.i. and washed 10 hours later for synchronous O/N rupture. The following day when parasites were ∼10 h.p.i., they were further synchronised by incubation in 5% w/v D-sorbitol to remove any remaining late-stage parasites. Cultures were then diluted to 1% parasitaemia/3% haematocrit and split across 5 x 5 mL cultures. To induce mislocalisation, 250 nM of Rapalog A/C heterodimeriser (Takara) was added to individual wells at the following time points – 10 h.p.i., 19 h.p.i., 31 h.p.i., and 40 h.p.i. 48-hours post-assay set-up, culture media was replaced and parasites were incubated for a further 24-hours.

### Assay analysis

Blood smears were made at each time point, fixed in 100% methanol, stained with 5% Giemsa and imaged using a Zeiss Axiolab 5 microscope.

Parasitaemia, life stage distribution and aberrant cell types were counted from end point Giemsa smears. Each culture condition was counted three times across separate days to ensure consistency.

### Immunofluorescence assays

Immunofluorescence assays were undertaken as described previously with modifications (Liffner *et al*, 2022). High parasitaemia (>5%) cultures were diluted to 0.3% haematocrit and incubated on 0.01% poly-L-lysine (Sigma-Aldrich) coated coverslips for 30 min at 37°C then washed with MilliQ H_2_O and remaining adhered parasites fixed with 4% paraformaldehyde (PFA) (Sigma-Aldrich) or 4% PFA/0.075% glutaraldehyde (Electron Microscopy Sciences) for 20 min at 37°C. Parasites from biotinylation assays were smeared directly onto glass slides and fixed in -20°C methanol for 30 min and stored at -80°C until staining.

Fixed parasites were permeabilised in 0.1% v/v Triton-X100 for 10 min, then incubated in 3% w/v bovine serum albumin (BSA)-PBS-T 0.05% blocking buffer for one hour at RT. Primary antibodies were diluted in 1% BSA-PBS-T 0.05% and added to coverslips O/N at 4°C to the following concentrations: 1:1000 rabbit anti-GFP (Genescript), 1:500 mouse anti-Exp2, 1:250 anti-streptavidin AF488 (Invitrogen). After removal of primary antibodies, coverslips/slides were washed three times in 1X PBS-T 0.05% wash buffer then incubated with DAPI (1:1000) and secondary antibodies (1:500 goat-anti-rabbit AF488, 1:500 goat-anti-mouse AF594) diluted in antibody diluent for 1-hour in the dark at RT. For PFA and PFA/glutaraldehyde-fixed coverslips, antibodies were post-fixed for 10 min RT then washed once with MilliQ and dehydrated in ascending concentrations of ethanol (70%, 90%, 100%) for 3 min each. Coverslips were then air-dried and mounted on glass slides with Prolong Gold or Prolong Diamond mounting medium and left to harden for 72-hours before imaging. Methanol fixed slides were air-dried, and Prolong Gold mounting media was used to affix coverslips.

All imaging was performed using a Zeiss LSM980 fluorescence microscope with a Plan-Apochromat 63x/1.4 oil DIC objective lens and 405 nm, 488 nm, and 594 nm laser lines and images were Airyscan processed.

### Live cell imaging

Parasites were diluted to 0.3% haematocrit and supplemented with 300 nM DAPI and for dimerisation assays, 250 nM of Rapalog was also added as appropriate. iBidi 15 μ-slide eight-well glass-bottom slides were used for imaging and coated with 0.01% poly-L-lysine, 200 μL of sample was added per well and incubated for 1-2 hours prior to imaging. Parasites were then imaged using a Nikon Ti2E fluorescent microscope using a Plan Apochromat 60x NA 1.4 oil immersion objective lens. Light sources were DAPI-5060C-NTE-ZERO, FITC-3540C-NTE-ZERO, and TxRED-4040C-NTE-ZERO.

### Image analysis

All microscopy images were analysed using Fiji/ImageJ software. Images were adjusted for brightness, contrast, and/or colour balance and cropped with scale bars added as appropriate.

### Culture and harvest of parasites for BioID experiments

PfAlba4^FKBP-GFP/mTID-FRB-mCh^ parasites were sorbitol synchronised and grown at ∼5% haematocrit in 125 mL volumes. Once parasitaemia reached 10-12% schizonts, 150 nM ML10 was added to ensure high synchronicity. For each replicate, media was removed from two 125 mL cultures and combined through resuspension in 60 mL of fresh biotin-free media then split between 2x 30 mL culture dishes and 5 mL set aside to serve as an untreated control for western blot validation. Both 30 mL cultures were supplemented with 150 µM of biotin and one was also treated with 250 nM of Rapalog (experimental sample). Cultures were then incubated at 37°C in a pre-warmed gas box under standard gas conditions for 30-mintues.

Parasites were then resuspended and washed twice in 1X PBS to remove ML10, excess biotin, and residual Albumax. Red blood cells were then lysed in 0.15% ice-cold saponin with 1X Protease inhibitor as described above.

### Enrichment of biotinylated peptides

Saponin pellets were resuspended in lysis buffer (2% SDS, 25 mM Tris-HCl at pH 8, 150 mM NaCl, 250 U/mL benzonase nuclease (Millipore), MilliQ H_2_O, 1X phosphatase inhibitor (Roche), 1X protease inhibitor (Cømplete, Roche) and sonicated at 50% Amp for 30-seconds until homogenous. Samples were diluted six times in -20°C acetone (Sigma) and incubated for 2-hours at -20°C to precipitate proteins, followed by 10-minute centrifugation at 4°C/13,000 x g. After air-drying, pellet was re-solubilised in 8 M urea (Sigma)/50 mM ammonium bicarbonate (ABC) (Sigma) and denatured at 38°C/650 rpm for 1-hour. Protein concentration for each sample was assessed by BCA assay as per the kit protocol and normalised across samples. Samples were then reduced and alkylated with 10 mM Bond-breaker Tris(2-carboxyethyl)phosphine TCEP (Thermofisher), 40 mM 2-chloroacetamide (CAA) (Sigma), then diluted 8 times with 50 mM ABC for a final urea concentration of 1 M and incubated at 55°C/650 rpm for 15 min. CaCl_2_ was added to a final concentration of 1 mM and incubated at 45°C/650 rpm for a further 15 min. Trypsin/Lys-C (Promega) was added to a 50:1 w/w ratio and samples incubated for 16-hours at 37°C/650 rpm.

To enrich for biotinylated peptides, 100 µl of streptavidin magnetic beads (Genscript, Chameleon Science) were prepared for each sample and washed in 1 ml of wash buffer (2 M urea, 25 mM Tris-HCl at pH 8, 150 mM NaCl). Trypsinised samples were clarified at 10,000 x g for 3 min and supernatant was used for bead re-suspension and incubated RT for 4-hours with gentle end-over-end rotation. A magnetic rack was used to remove the supernatant, and beads were washed twice in 1 ml of wash buffer and once in mass spec grade H_2_O. Biotinylated peptides were released from beads by 5-minute incubations at 60°C in elution buffer (80% acetonitrile (ACN) (Sigma), 0.2% trifluoroacetic acid (TFA) (Sigma), 0.1% formic acid (FA) (Thermofisher)), beads were magnetised and the peptide-containing supernatant removed to a new tube, this was repeated five times. Samples were then dried by vacuum centrifugation to remove ACN and desalted using C18 stage tips.

### GFP coimmunoprecipitation assays

Synchronised parasites at 10-12% parasitaemia/3% haematocrit were harvested by saponin lysis as described above. Lysates were then treated with 250 μl of lysis buffer (10 mM Tris, 5 mM EDTA, 20 mM beta-glycerolphosphate pH 7.5, 1% (v/v) Nonident-40 (NP-40) and Complete Protease Inhibitor Cocktail (Roche)) and incubated for 15 min at 4°C then centrifuged for 3 min/14,000 x g. Supernatant was collected then pellets were further solubilised in 200 μl of wash buffer (10 mM Tris, 5 mM EDTA, 20 mM beta-glycerolphosphate pH 7.5) and probe sonication (10 amp/15-seconds/three times) on ice. Samples were pelleted again and supernatant pooled with the NP-40 soluble fraction and sample volumes were adjusted to reach a final NP-40 concentration of 0.1%.

GFP trap® resins were washed three times in 100 μl of wash buffer and then resuspended with parasite lysates. Samples were incubated on a rotor for 2-hours at 4°C then pelleted at 600 x g for 2 min and washed three times with wash buffer. Resins were resuspended in 2X reducing sample buffer and separated by SDS-PAGE. Gels were then stained with Instant Blue stain (Expedion) for 30 min, gently rocking, then destained O/N in water. The region of gel where sample was resolved was cut from the gel for mass spectrometric analysis.

Excised gel pieces were washed in 10X gel volume of 100 mM ABC and further destained by addition of 10X gel volume of 50% ACN in 50 mM ABC and gentle agitation for 20 min. Supernatants were then discarded and the above process was repeated to achieve complete destaining.

De-stained gel pieces were reduced using dithiothreitol (DTT) at a final concentration of 5 mM in 100 mM ABC and incubated for 30 min at 65°C. Proteins were then alkylated using CAA at 11 mM final concentration and incubated for 30 min RT in the dark. Supernatant was then discarded and gels washed once with 500 μl of 50% ACN/50 mM ABC and twice in 10X gel volume of ACN for 15 min at RT on roller for gel dehydration. Following, 200 μl of trypsin (Promega) was added (12.5 ng/μl) and incubated O/N at 37°C.

The following day digestion was quenched with 0.5X gel volume of extraction buffer (50% ACN, 5% FA) and the digested peptides were extracted and transferred to new tubes and dried by speed-vac (VWR International PTY LTD). Samples were then resuspended in 20 μl of 2% ACN with 0.1% FA for LC-MS/MS analysis.

### Mass spectrometric analysis of BioID and CoIP samples

Protein concentration was normalised across samples for both experiments and were analysed using Vanquish Neo nano LC system with an Orbitrap Astral mass spectrometer with NanoSpray Flex source (Thermo Scientific). For BioID experiments a 50 cm µPAC^TM^ HPLC column was used (Thermo Scientific) with a 60 min acquisition time and for CoIP experiments, Aurora Ultimate 25x75 C18 UHPLC columns (IonOpticks) with a 20 min acquisition time.

### Analysis of BioID data

BioID raw mass spectrometry files were processed using PEAKS Studio 13 (Bioinformatics Solutions). Data refinement was performed with default instrument-specific parameters. Database searching was conducted against an in-house spectral library of *P. falciparum* proteins and the [Homo Sapiens SwissProt (June 2023), (UP000001450, release version 2016_04)] appended with a library of common laboratory contaminants (Siddiqui *et al*, 2022a; Siddiqui *et al*, 2022b).

The search parameters were as default except Biotinylation (K, +226.0776 Da) was additionally included as a variable modification, allowing a maximum of three variable modifications per peptide. A false discovery rate (FDR) threshold of 1.0% was strictly applied at both the peptide-spectrum match and protein levels.

Further analysis of BioID data was undertaken directly at the peptide level. First, peptides were filtered to include only those with a +226.0776 biotin modification. Intensity values of these peptides were then log10 transformed, missing values were assigned as ‘0’, and a p-value generated by students t-test. The harmonic mean of individual biotinylated peptide p-values was then calculated to obtain protein level p-values. Total biotinylated peptides for each protein were used to calculate foldchange (#peptides in +Biotin/+Rapalog samples / #peptides in +Biotin only control samples) then log2 transformed. Where there were no biotinylated peptides detected in the +Biotin controls, the foldchange was calculated as the sum of peptides in +Biotin/+Rapalog samples and log2 transformed. Proteins were considered significantly enriched if they had a biotinylated peptide log2 fold change ≥2 in the +Biotin/+Rapalog samples and A) were present in all +Biotin/+Rapalog replicates or B) were present in two or more +Biotin/+Rapalog replicates with at least two peptides detected in two replicates.

### Analysis of coimmunoprecipitation data

Protein identifications and associated label-free quantification values were obtained from raw data files using MaxQuant version 1.6.17 with Plasmodium protein annotations retrieved from PlasmoDB and human protein annotations from [Homo Sapiens SwissProt (June 2023), (UP000001450, release version 2016_04)]. The following analysis parameters were used: 1% FDR cutoff value and the following modifications - carbomidomethylation (fixed), Oxidation @M and Acetylation @ protein N-terminus (variable).

Data were then analysed with the in-house LFQ analyst software. Briefly, all contaminant proteins, reverse sequences and proteins identified ‘only by site’ were excluded as well as proteins where only a single peptide was identified or consistently quantified. Samples were grouped by condition (GFP or WT). Missing not at random (MNAR) imputation of missing values was used and differentially expressed proteins were identified by pairwise comparisons of GFP vs WT samples. The Benjamini-Hochberg method was used to calculate the adjusted p-value of each protein and cut-off values of ≤ 0.05 adjusted p-value and log2 foldchange ≥ 2 were used to identify significant proteins in the data.

### PfAlba4 protein immunoprecipitation and RIP-seq

Schizont-stage parasites from the PfAlba4 FKBP-GFP transgenic line were harvested in three independent biological replicate experiments for RNA immunoprecipitation sequencing (RIP-seq). Each replicate comprised approximately 10–15 × 10⁶ tightly synchronised late-schizont parasites. Infected erythrocytes were lysed with 0.05% (w/v) saponin in ice-cold 1× PBS supplemented with protease inhibitor for 2–3 min on ice. Parasites were collected by centrifugation at 16,000 × g for 5 min at 4 °C and washed three times with ice-cold 1× PBS containing protease inhibitor.

Washed parasite pellets were resuspended in 10 pellet volumes of extraction buffer containing 1% NP-40, 150 mM HEPES and 150 mM NaCl (pH 7.0), supplemented with protease inhibitor. Samples were sonicated on ice for 30 s at 20% amplitude using 1-s on/1-s off pulse cycles, incubated overnight at 4 °C with rotation, and clarified by centrifugation at 16,000 × g for 20 min at 4 °C.

Ten per cent of each clarified lysate was retained as the unenriched total-RNA input control. The remaining lysate was incubated with 20 μL anti-GFP magnetic beads for 2-hours at 4 °C with gentle rotation. Beads were collected using a magnetic rack and washed sequentially five times with 1 mL extraction buffer, five times with 1 mL 0.5× extraction buffer and five times with 1 mL nuclease-free water. Washed bead-bound complexes were subjected to proteinase K digestion before RNA extraction.

For input samples, 100 μL of the reserved lysate was mixed with 1 mL TRIzol reagent. Immunoprecipitated RNA was extracted by directly resuspending the washed beads in 1 mL TRIzol following proteinase K digestion. Samples were mixed with 200 μL chloroform, vortexed briefly and centrifuged at 13,000 × g for 15 min at 4 °C. RNA was precipitated from the aqueous phase by addition of 500 μL ice-cold isopropanol and incubation for 10 min at room temperature, followed by centrifugation at 12,000 × g for 10 min at 4 °C. RNA pellets were washed once with 1 mL ice-cold 70% ethanol, centrifuged at 7,500 × g for 5 min at 4 °C, air-dried and resuspended in 50 μL nuclease-free water.

RNA was treated with TURBO DNase according to the manufacturer’s protocol. Briefly, 0.1 volumes of 10× TURBO DNase buffer and 1 μL TURBO DNase were added to each sample, followed by incubation at 37 °C for 20–30 min. DNase inactivation reagent was subsequently added, samples were incubated for 5 min at room temperature and centrifuged at 10,000 × g for 1.5 min, and the RNA-containing supernatant was transferred to a fresh tube. RNA concentration was determined using a Qubit fluorometer with the RNA High Sensitivity assay, and RNA integrity was assessed using an Agilent TapeStation.

### Library preparation, sequencing and read processing

Input and PfAlba4 RIP RNA libraries were prepared using the GenScript Ultra-Low Input mRNA Non-Directional Library Prep Kit and sequenced as 150-bp paired-end reads on an Illumina MiniSeq. Raw read quality was assessed using FastQC. Reads were trimmed using Trimmomatic v0.39 with a four-nucleotide sliding window, a minimum mean quality score of 20, removal of three nucleotides from both read ends and a minimum retained length of 50 nt. Trimmed reads were aligned to the *Plasmodium falciparum* 3D7 reference genome (PlasmoDB release 68) using Subread, and transcript-level counts were obtained using featureCounts with the corresponding PlasmoDB annotation.

### RIP-seq enrichment analysis and target identification

For enrichment analysis, featureCounts values were combined with transcript-length information and converted independently for each library to transcripts per million (TPM). Transcripts were retained when >10 raw reads were detected in at least two of three RIP libraries and independently in at least two of three input libraries. For each retained transcript, RIP and input abundance were summarised as the median TPM across the three biological replicates, and enrichment was calculated as the RIP-to-input median TPM ratio after addition of a 1-TPM pseudocount. Enrichment values were subsequently log2-transformed.

The distribution of log2(RIP/Input) enrichment values was fitted using a two-component Gaussian mixture model. The components with the lower and higher fitted means were designated the lower-enrichment and RIP-enriched statistical populations, respectively. For each transcript, the posterior probability of membership in the lower-enrichment component was treated as a local model-based false-discovery probability. Transcripts were ranked from highest to lowest enrichment, and a GMM-derived cumulative false discovery rate (FDR) was calculated as the running mean of these probabilities.

Primary PfAlba4-associated transcripts were defined by ≥2.5-fold RIP/Input enrichment and GMM-derived cumulative FDR ≤0.05. A posterior probability ≥0.95 for assignment to the RIP-enriched component, equivalent to a natural-log posterior odds score (LOD) ≥2.94, was retained as an additional confidence measure but was not used as an independent primary target criterion. A stringent high-confidence subset was defined by ≥5-fold enrichment and GMM-derived cumulative FDR ≤0.01.

### 3′UTR motif analysis

To identify putative cis-regulatory elements associated with PfAlba4 target recognition, 3′UTR sequences were extracted from the *P. falciparum* 3D7 reference genome using PlasmoDB release 68 annotation. PfAlba4 target 3′UTRs of at least 50 nt were retained for analysis. Non-target 3′UTRs were obtained from transcripts evaluated in the RIP-seq experiment that were not significantly enriched by either PfAlba2 or PfAlba4 and were used as the discriminative background set, to avoid signal contamination in the negative set.

XSTREME (MEME Suite v5.5) was run in discriminative RNA mode using motif widths of 6–15 nt, with the non-target sequences used to construct the background model. Motif enrichment was assessed using SEA with Fisher’s exact test, with statistical significance reported as E-values and FDR-adjusted q-values. Discovered motifs were compared with experimentally characterised RNA-binding protein motifs in the Ray2013 RNAcompete database using Tomtom (Ray *et al*., 2013). FIMO was used to identify individual PfAlba4-associated 3′UTRs containing selected enriched motifs.

### Functional enrichment analysis

Functional enrichment analysis of the primary PfAlba4-associated transcript set was performed using STRING v12.0. *P. falciparum* 3D7 identifiers were mapped using the default STRING resolver, with duplicate mappings collapsed and unmapped identifiers excluded. The organism-wide *P. falciparum* 3D7 proteome was used as the reference background.

Gene Ontology Biological Process, Molecular Function and Cellular Component categories, together with KEGG and Reactome pathways, were assessed using the STRING enrichment framework. Multiple-testing correction was performed using the Benjamini–Hochberg procedure, with FDR <0.05 considered significant. Enrichment magnitude was represented by STRING Strength, defined as log10(observed/expected), and visualised using bubble or lollipop plots with point size representing the number of genes contributing to each enriched term.

### Protein–protein interaction network analysis

Protein–protein interaction networks were generated using STRING v12.0 with the default evidence channels and a medium-confidence interaction threshold of combined score ≥0.4. Disconnected nodes were excluded from network visualisation. Network modules were identified using the Markov Cluster Algorithm (MCL) with an inflation parameter of 3. Node size represented network degree, edge thickness represented the STRING combined interaction score, and module membership reflected the corresponding MCL cluster.

### Comparison with stage-specific proteomes

To assess stage-specific protein detection of PfAlba4-associated transcripts, the 692 primary PfAlba4 targets were compared with schizont and wild-type Day 3 gametocyte proteomic datasets (Chisholm *et al*., 2026; Marapana *et al*., 2026). Comparisons were performed at the PF3D7 gene level after removal of transcript-version suffixes.

A target was considered detected in the schizont proteome when its corresponding PF3D7 identifier was present in the schizont proteomic dataset. For the Day 3 gametocyte dataset, proteins were considered detected when classified as identified in wild-type Day 3 gametocytes. Where quantitative Day 3 measurements were available, protein abundance was represented by the mean normalised signal across the five wild-type Day 3 samples.

PfAlba4-associated transcripts were classified as detected in both proteomes, schizont only, Day 3 gametocyte only, or neither dataset. This analysis was used to determine the proportion of PfAlba4-associated transcripts with corresponding protein-level evidence at the schizont stage and the proportion of schizont-undetected targets subsequently detected during early gametocyte development.

### Enrichment analysis

Gene Ontology enrichment analysis was performed using ShinyGo version 0.85.1 with default parameters (0.05 FDR cut-off, minimum pathway size = 2, redundancy removed, # of pathways to show = 20). K-means clustering was performed in StringDB.

### Statistical analysis

All statistical analysis and graphs were generated using GraphPad PRISM 11 (GraphPad Software Inc.). Statistical tests and experimental information are indicated in figure legends as appropriate.

## Data Availability

Proteomics and RIP-Seq data will be made available on an online repository upon acceptance of the manuscript. All other data and reagents are included in the manuscript or available through contacting the corresponding author.

## Acknowledgements

D.W.W. and D.J.C. are ARC Future Fellows. A.R.J. is an NHMRC Investigator Fellow. This work was supported by an ARC Discovery Project (DP250104773). M. E. Z. was supported by an ARC FT24 scholarship.

